# NLRP3 inhibition maintains microglia architecture and enhances behavioral recovery after traumatic brain injury

**DOI:** 10.1101/2025.06.25.660862

**Authors:** Sergio Castro-Gomez, Ana Vieira-Saecker, Juan Ignacio Muñoz-Manco, Tao Li, Ida Kulińska, Felix D Weiss, Felix Meissner, Stephanie Schwartz, Valentin Stein, Eicke Latz, Michael T. Heneka

**Author notes:** Correspondence (S.C-G.), (M.T.H.).

## Abstract

NLRP3 plays an essential role in secondary neuroinflammatory damage following traumatic brain injury (TBI). However, the specific mechanisms mediating NLRP3’s effects in TBI remain poorly understood, and it is unknown whether its pharmacological inhibition with oral compounds during initial phases confers long-term protection. In this study, we investigated the role of the NLRP3 inflammasome pathway in TBI-induced neuroinflammation and long-term neurobehavioral impairment, as well as the impact of its pharmacological inhibition. Following controlled cortical impact (CCI), most NLRP3 inflammasome-related proteins and key inflammatory markers were elevated during the first week post-injury. Genetic *Nlrp3* deletion and treatment with oral NLRP3-specific inhibitors preserved microglial homeostatic architecture, reduced ASC aggregation, and enhanced neurological and cognitive recovery after CCI. Repeated intravital imaging confirmed that NLRP3 inhibition prevents microglial activation post-TBI. These findings suggest NLRP3 inflammasome targeting represents a viable translational strategy for clinical trials and may mitigate long-term neurocognitive decline following TBI.

## INTRODUCTION

Microglia are key players in the innate immune responses of the brain and act as first responders to neuroinflammatory stimuli^1^. In parallel, activation of the Nucleotide-binding oligomerization domain-, leucine-rich repeat-, and pyrin domain-containing 3 (NLRP3) inflammasome is a pivotal event in cerebral innate immunity^2^. The NLRP3 inflammasome is composed of the sensor molecule NLRP3, the adaptor protein Apoptosis-associated Apeck-like protein Containing a CARD (ASC), and the effector molecule caspase-1 (CASP1)^3^. Upon stimulation, NLRP3 undergoes conformational changes that relieve its autoinhibitory state, enabling oligomerization and the formation of helical fibrillar assemblies with ASC. ASC filaments serve as a molecular platform to recruit pro-caspase-1 (pro-CASP1), facilitating its cleavage and subsequent activation via proximity-induced mechanisms. Cleaved caspase-1 (CASP1) then processes the pro-forms of interleukin-1β (IL-1β) and IL-18, and mediates the proteolytic activation of gasdermin D (GSDMD). GSDMD forms pores in the cellular membrane that are crucial for the release of proinflammatory mediators and for inducing pyroptosis, a pro-inflammatory form of cell death triggered by sustained inflammasome activation^4^. The NLRP3 inflammasome can be activated by a broad range of stimuli following direct mechanical brain injury, including altered ion homeostasis, excessive glutamate^5^, release of ATP^6^, calcium dysregulation^7^, potassium efflux^8^, and the generation of reactive oxygen species^9^.

Traumatic brain injury (TBI) is a leading cause of death and disability in the economically active population, with limited medical options to prevent long-term neurocognitive deficits^10^. TBI is also recognized as the most significant non-genetic, non-age-related risk factor for developing dementia, including Alzheimer’s Disease (AD)^11^. Tissue damage from TBI is classified as either primary or secondary. Primary TBI refers to the immediate mechanical insult and its direct consequences, such as hematoma or hypoxia. Secondary TBI arises from subsequent injury cascades, including glutamatergic excitotoxicity, calcium overload, vascular dysfunction, and neuroinflammation. Secondary damage can persist for months to years and contributes to chronic cognitive dysfunction and dementia^12^. In humans and murine models, increased production of IL-1β, characterizes the initial immune response after TBI, underscoring the central role of this cytokine and inflammasome activation in TBI-induced inflammation^13,14^. Levels of NLRP3^15^ and other inflammasome-related proteins (e.g., ASC, CASP1, IL-18) rise significantly in serum and cerebrospinal fluid after TBI, and higher concentrations correlate with poorer clinical outcomes^16^.

Previous studies have shown that *Nlrp3* deficiency in mice confers protection against acute neurocognitive impairment following closed head injury (CHI)^17,18^. Pharmacological inhibition of NLRP3 using agents such as BAY11-7082^17^, JC124^19^ or MCC950^20–22^ has demonstrated neuroprotective and anti-inflammatory effects in rodent TBI models. However, some of the reports are controversial. For instance, it was previously described that *Nlrp3* knockout (*Nlrp3^-/-^*) mice or those treated with MCC950 prior to TBI may experience greater disruption of the blood-brain barrier (BBB) compared to control animals. Thus, the precise mechanism of NLRP3 activation in post-TBI cerebral dysfunction, particularly in later phases, is still an open question. It also remains unclear whether treatment with NLRP3 antagonists in the initial post-TBI phases can prevent long-term neurocognitive impairment.

In this study, we performed biochemical and histological characterization of *Nlrp3^-/-^* mice subjected to CCI as a model for moderate TBI model, with a particular focus on later stages (up to 30 days post-injury) and using microglial morphology as a primary neuroimmunological readout. Additionally, we investigated whether pharmacological inhibition of NLRP3 with recently developed oral compounds (NP3-253, NP3-361) during the acute phase (first 10 days post-injury) could prevent long-term neurocognitive impairment, and microglial reactivity *in vivo* using repeated intravital two photon laser microscopy (2PLM). We found that NLRP3 inflammasome-related proteins and key inflammatory markers are upregulated especially the first 7 days post injury. Microglia from *Nlrp3^-/-^* or NHY361-treated mice retained a homeostatic-like morphology. Furthermore, injured *Nlrp3^-/-^* mice or those treated with oral NLRP3 inhibitors showed faster neurological recovery and better performance in neurobehavioral tests during the initial phases of injury compared to vehicle-treated wild-type controls.

## RESULTS

In this study, we utilized CCI as common and widely used procedure to model TBI in rodents. CCI is induced using an electromagnetic device that drives a rigid impactor onto the exposed, intact dura, resulting in cortical tissue loss, axonal injury, concussion, and BBB disruption^23^. Initially, we performed CCI on 16 mice (8 wild-type [WT] and 8 *Nlrp3^-/-^*) and estimated the percentage of tissue loss in brain sections after Nissl staining at 3- and 30 days post-impact (dpi) (Figure 1A-B). We observed a significant genotype-dependent difference in tissue loss at both points (3 dpi: WT, 12.12 ± 0.69% vs. *Nlrp3^-/-^*, 6.79 ± 1.17%; 30 dpi: WT, 11.24 ± 2.81% vs. *Nlrp3^-/-^*, 5.49 ± 2.60%). Interestingly, TUNEL staining of perilesional hippocampal tissue at 3 dpi revealed no difference in the number of apoptotic cells between the two genotypes (Figure S1). This suggests that a form of cell death other than apoptosis contributes to the NLRP3-mediated tissue loss observed in this model.

**Figure 1.**
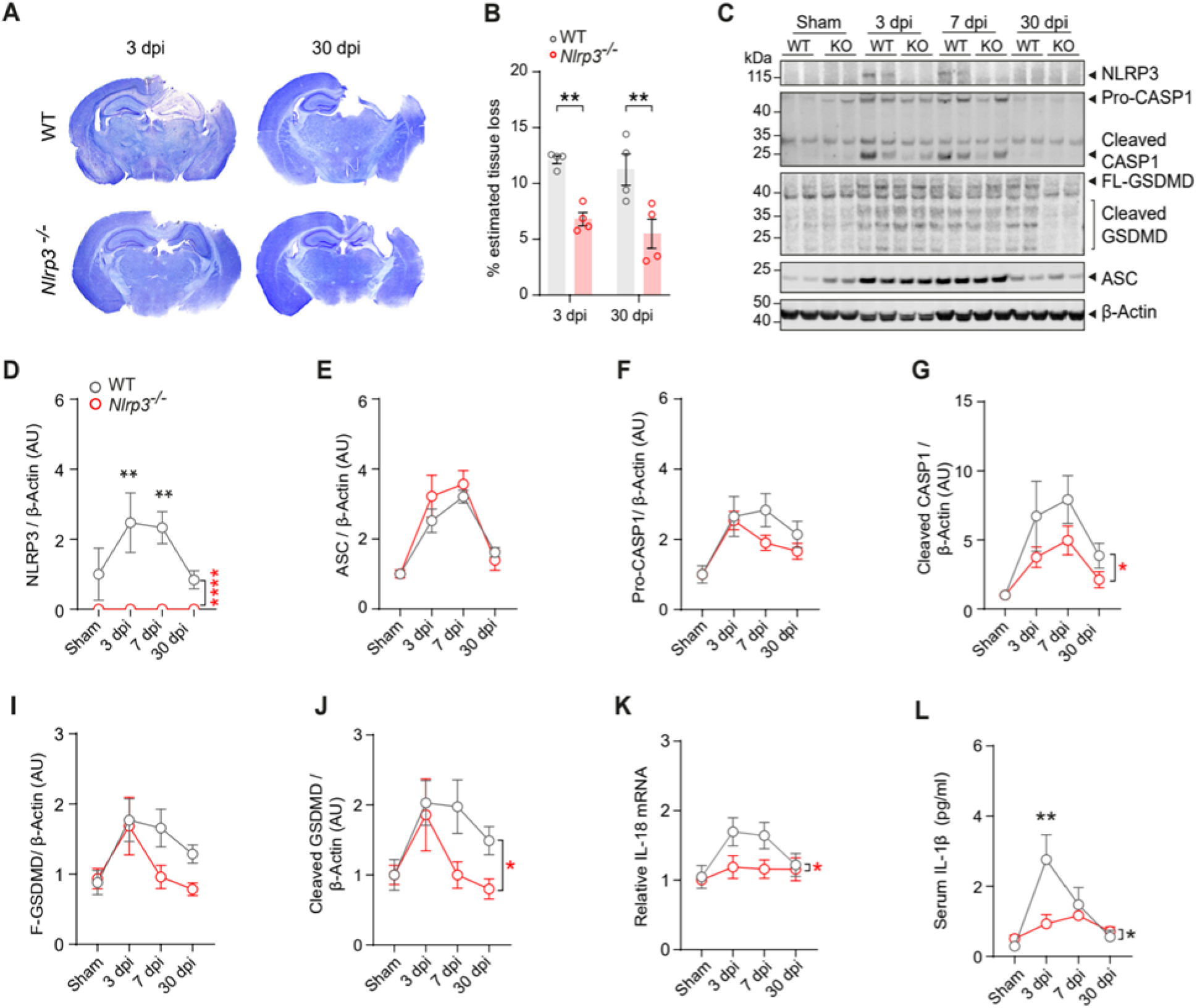
NLRP3 inflammasome-related proteins are consistently upregulated after brain injury. (A) Representative Nissl-stained brain sections from WT and *Nlrp3^-/-^* at 3- and 30-days post-CCI (dpi). (B) *Nlrp3^-/-^* mice exhibit significantly reduced tissue loss at 3 and 30 days following CCI compared to injured WT controls. n = 4 mice/genotype/timepoint; individual data points represent biological replicates. Two-way ANOVA (factors: genotype × time). (C) Representative Immunoblot and (D-J) densitometric analysis of NLRP3, ASC, CAPS1 and GSDMD in perilesional cortex at 3-, 7- and 30 days dpi. (D) NLRP3 increases 3- and 7-days post injury. n = 5 mice/genotype/timepoint. Two-Way ANOVA (E) ASC expression rises temporally in WT and *Nlrp3^-/-^* mice. n = 5 mice/genotype/timepoint. (F) Pro-CASP1 elevation at 3 and 7 dpi. n = 5 mice/genotype/timepoint. Two-Way ANOVA (G) Reduced cleaved Casp1 in *Nlrp3^-/-^* mice across timepoints. n = 5 mice/genotype/timepoint. Two-Way ANOVA. (H) GSDMD upregulation peaks between 3–7 dpi. n = 4-5 mice/genotype/timepoint. Two-Way ANOVA. (I) Attenuated cleaved GSDMD in *Nlrp3^-/-^* mice post-CCI. n = 4-5 mice/genotype/timepoint. Two-Way ANOVA. (J) Perilesional IL-18 mRNA levels are suppressed over time following CCI in *Nlrp3^-/-^* mice compared to WT controls. n = 5 mice/genotype/timepoint. Two-Way ANOVA. (K) Nlrp3^-/-^ mice displayed suppressed IL-1β response in serum over time following CCI. n = 5 mice/genotype/timepoint. Two-Way ANOVA. All data are presented as mean ± SEM. Significance is indicated in red for genotype effects and in black for interaction effects. All post hoc analyses were performed using the Bonferroni test. *p < 0.05, **p < 0.01, ***p < 0.001, ****p < 0.0001.

### NLRP3 inflammasome-related proteins and other key inflammatory mediators are upregulated following moderate TBI

To investigate NLRP3 inflammasome-related proteins following CCI, we first dissected perilesional cortices from WT and *Nlrp3^-/-^* mice at three different time points (3-, 7-, and 30 dpi) and performed immunoblot analyses for NLRP3, ASC, CASP1, and GSDMD (Figure 1C). Densitometric analysis revealed a significant upregulation of these proteins, peaking between 3 and 7 dpi. By 30 dpi, the levels of NLRP3 and ASC nearly return to baseline (Figure 1D-E), whereas the expression and cleavage of CASP1 and GSDMD remained elevated (Figure 1F-I). Consistent with the essential role of NLRP3 in neuroinflammatory responses following CCI, the constitutive absence of *Nlrp3* gene resulted in a significant genotype-dependent reduction of cleaved CASP1 (Figure 1G) and cleaved GSDMD (Figure 1J) over time. Cleaved CASP1 represents the inflammasome’s primary effector protein, which processes some mediator such GSDMD, IL-1β and IL-18^2,4^.

To identify the effect of *Nlrp3* deletion on additional neuroinflammatory mediators, we performed multiplex ELISA assays in perilesional cortices and serum, as well as quantitative PCR (qPCR) for proinflammatory markers in perilesional cortices from WT and *Nlrp3^-/-^* mice at 3-, 7-, and 30 dpi following CCI (Figure S2A-S). Similar to NLRP3 inflammasome-related proteins, IL-1β, IL-6, IL-10, TNF-α, IFN-γ, CXCL1, along with *Il-18*, *C1q*, and *C3* mRNA levels were upregulated in perilesional cortices, peaking between 3 and 7 dpi (Figure S2A-S). This pattern confirms a pronounced acute and subacute neuroinflammatory response within the first week after TBI. While most of our measurements show no genotype-specific differences, *Nlrp3^-/-^* mice exhibited significantly impaired *Il-18* mRNA upregulation in perilesional tissue (Figure 1J) and an attenuated serum IL-1β response (Figure 1K). These findings indicate not only local, tissue-specific activation of the NLRP3 inflammasome, but also sustained, multisystemic NLRP3-dependent inflammation following CCI, which can remain detectable even 30 days post-injury.

### Differential distribution and aggregation of ASC in *Nlrp3^-/-^* brains after moderate injury

To gain a deeper understanding about the protective mechanisms from *Nlrp3* deficiency, we performed immunocytochemical analysis of the inflammasome adaptor protein ASC as proxy for inflammasome activation and pyroptosis initiation, and of the Allograft inflammatory factor 1 (Iba-1) to tract morphological changes in microglia as measurement of cellular neuroinflammation. Using a monoclonal knockout-validated antibody against ASC^24^, we detected and reconstructed ASC aggregates at 3- and 30 days post injury in WT and *Nlrp3^-/-^* mice subjected to CCI (Figure 2A). Upon activation, NLRP3 interacts with the adaptor protein ASC leading to its aggregation in filaments and other macromolecular aggregates such as ASC specks. ASC aggregates are considered a sing of inflammasome activation and pyroptosis^25^. Notably, *Nlrp3^-/-^* mice show significant lower amount of ASC aggregates at 3- and 30 days post-TBI (Figure 2B). Furthermore, *Nlrp3^-/-^* microglia presented significant less intracellular ASC aggregates at 30 days post CCI (Figure 2C). Additionally, we estimated the volume of ASC aggregates after three-dimensional reconstruction from confocal laser Z-Stack images. Similarly, ASC aggregates in *Nlrp3^-/-^*brains appear with significantly lower volumes at 3 days post injury (Figure 2D), especially those found within microglia cells (Figure 2E). These findings suggest that NLPR3 contribute to the normal distribution and aggregation of the adaptor protein ASC and subsequent processing of proinflammatory mediator after traumatic lesions.

**Figure 2.**
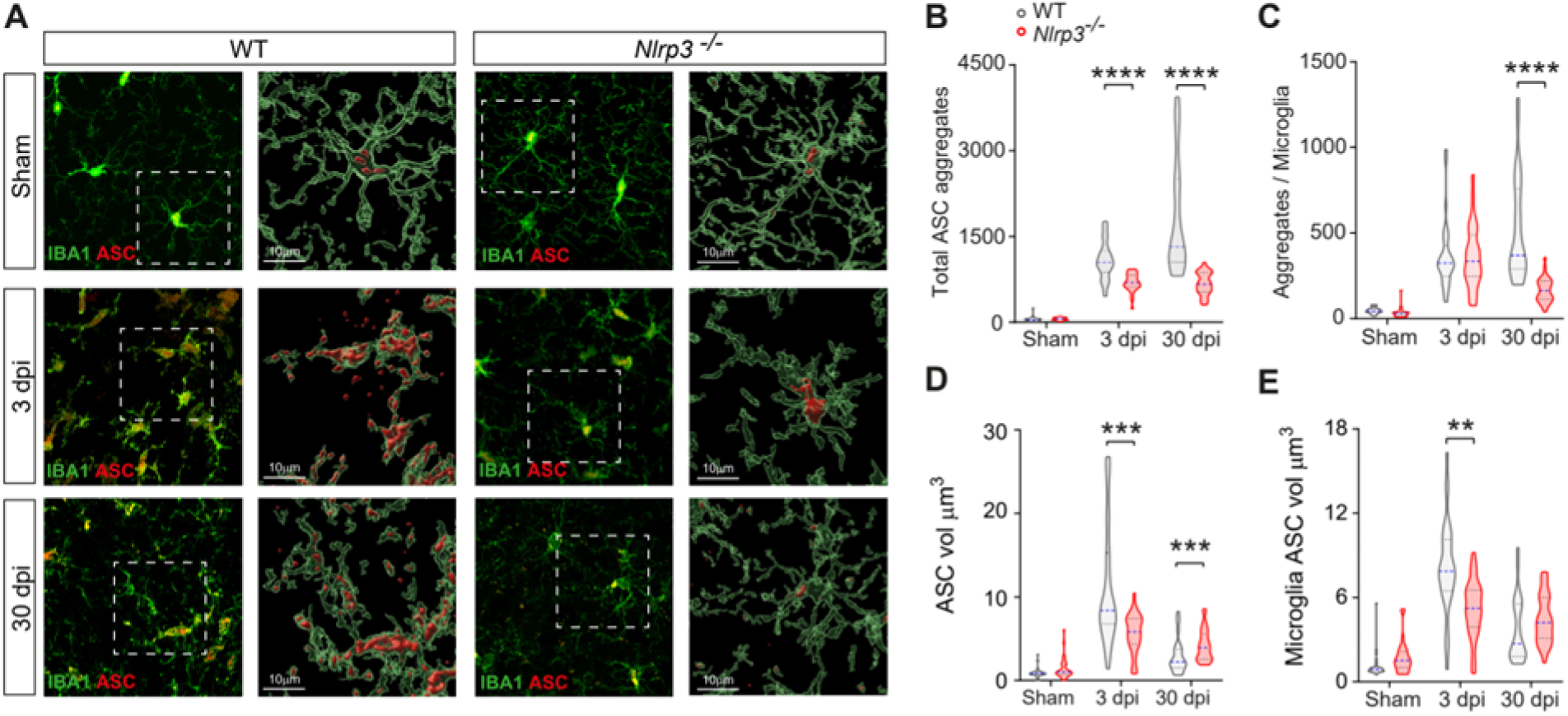
NLRP3 contributes to the aggregation and distribution of ASC after TBI. (A) Representative immunohistochemical images and surface 3D reconstructions of ASC and microglia at 3 and 30 days post-TBI. (B) *Nlrp3^-/-^* mice exhibit significantly fewer ASC aggregates at 3 and 30 days post-TBI. (C) *Nlrp3^-/-^* mice show reduced ASC aggregates per microglia at 30 days post-TBI. (D) Total volume of ASC aggregates is significantly reduced in *Nlrp3^-/-^* mice following CCI. (E) Volume of ASC aggregates within microglia is significantly lower in *Nlrp3^-/-^* mice 3 days post-injury. Truncated violin plots display the estimated data distribution, bounded by minimum and maximum observed values, with horizontal caps. Central and quartile lines indicate the median (purple dashed line) and interquartile range (black dashed lines). n = 7–8 sections per mouse from 4 mice/genotype/timepoint. ANOVA of Aligned Rank Transformed Data. All post hoc analyses were performed using the Tukey test. Significance is indicated as *p < 0.05, **p < 0.01, ***p < 0.001, ****p < 0.0001.

### NLRP3 is essential for morphological responses of microglia after brain injury

Spatiotemporal changes in microglial morphological phenotypes after trauma have been proposed as one of the best indicators of altered physiology in the lesioned tissue^26^. Using immunofluorescent staining against Iba-1, laser confocal microscopy and a previously validated MATLab based algorithm^27^, we reconstructed and analyzed fine morphological responses of microglia at 3- and 30 days following injury in WT and *Nlrp3^-/-^* mice (Figure 3A). *Nlrp3* genetic deficiency significantly prevents loss in microglial territorial volume (Figure 3B) with subsequent preserved branch length at 3 days post-TBI (Figure 3E). Furthermore, *Nlrp3^-/-^*microglia displayed significantly lower microglial cell volumes (Figure 3C) and higher number of endpoints (Figure 3D), higher number of branch points (Figure 3F) and significantly increased ramification index at 3 and 30 days post-TBI (Figure 3G) in comparison to WT-injured microglia. These observations suggest a diminished reactivity of *Nlrp3^-/-^* microglia with a resilient phenotype, possibly contributing to the significant smaller lesion sizes.

**Figure 3.**
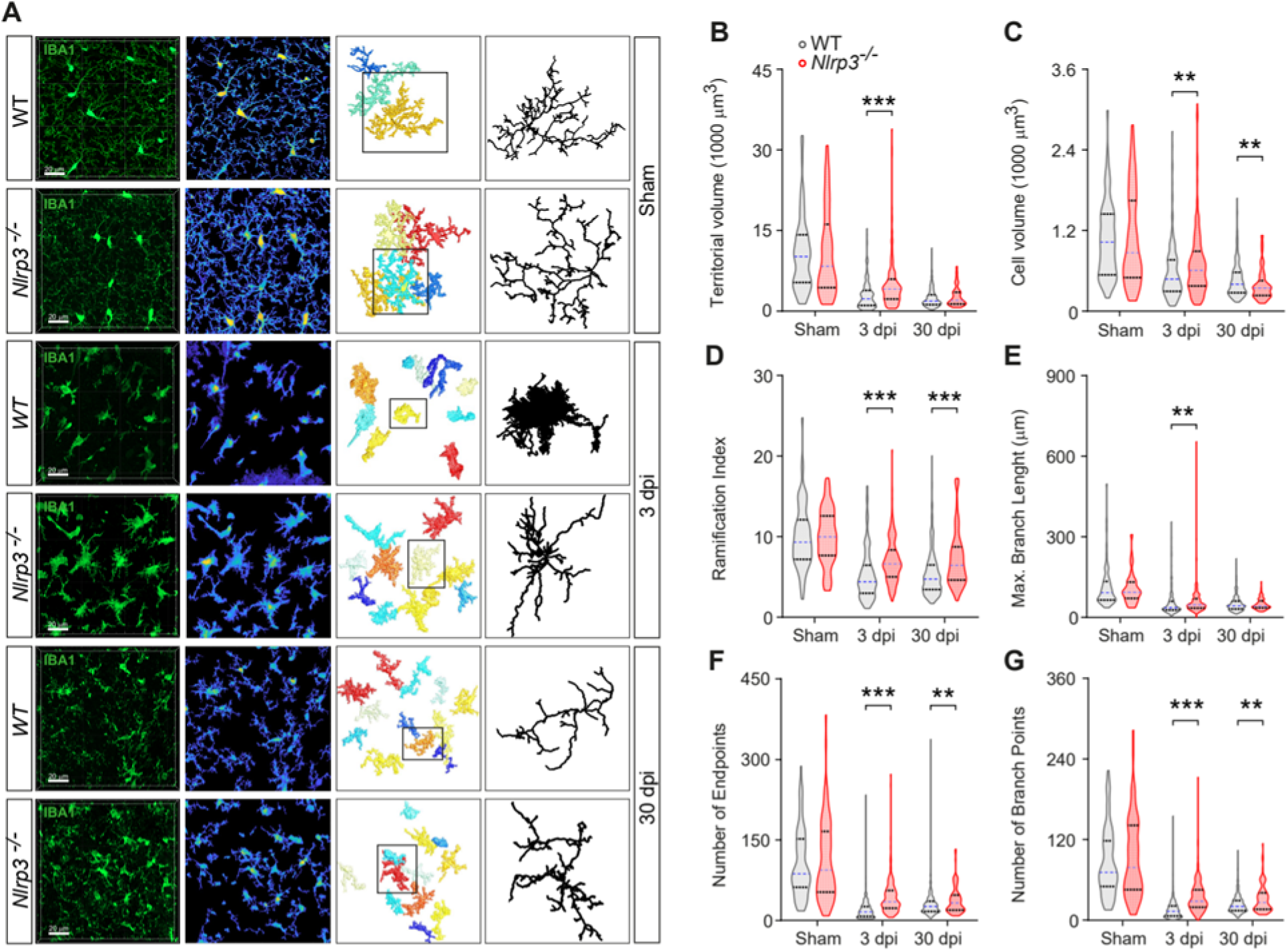
NLRP3 contributes to microglia morphological responses after brain injury. (A) Representative maximal intensity projections of confocal Z-stack images from Iba1-immunostained brain sections of WT and *Nlrp3^-/-^* mice at 3 and 30 days post-CCI. Shown are thresholded images, 3D reconstructions, and skeletonized examples of individual microglia processed and analyzed using 3DMorph.^27^. (B) *Nlrp3* genetic deficiency prevents TBI-induced reductions in territorial volume. (C) N*lrp3^-/-^* microglia display increased total cell volume at 3 days, but a significant reduction in somata volume at 30 days post-TBI. (D) Microglia from *Nlrp3^-/-^* mice exhibit a significantly higher ramification index at both 3 and 30 days post-TBI. (E) Maximal branch length is significantly greater in *Nlrp3^-/-^* microglia at 3 days post-TBI. (F) *Nlrp3^-/-^* microglia show a significantly higher number of endpoints at 3 and 30 days following CCI. (G) Number of branch points in *Nlrp3^-/-^* microglia at 3 and 30 days post-CCI. Truncated violin plots display the estimated data distribution, bounded by minimum and maximum observed values, with horizontal caps. The median (purple dashed line) and interquartile range (black dashed lines) are indicated. n = 74–320 cells from 7–8 sections per mouse, 4 mice per genotype/group. Statistical analysis was performed using ANOVA of Aligned Rank Transformed Data, with Tukey’s post hoc test. *p < 0.05, **p < 0.01, ***p < 0.001, ****p < 0.0001.

### NLRP3 contributes to acute and long-term behavioral phenotypes after CCI

TBI is not only manifested by motor and neurological deficits in the acute phase but with long-term neurobehavioral deficits in cognitive domains such as anxiety, executive functions, short- and long-term memory. To investigate, neurobehavioral phenotypes of *Nlrp3^-/-^* mice after moderate TBI, we performed a battery of behavioral tests that included the revised Neurobehavioral Severity Scale (NSS-R), Elevated O Maze (EOM), Open Field (OF), Novel Object Location Recognition task (NOLR) and Morris Water Maze (MWM) (Figure 4A and Figure 4B). As expected from our previous lesion volume analysis (Figure 1A), *Nlrp3^-/-^* mice showed a significantly faster recovery of neurological functions when evaluated with the NSS-R over time compared to injured WT controls (Figure 4B). In the EOM at 8 dpi, *Nlrp3*^-/-^ mice presented significantly longer exploration on the closed sectors, reflecting a conserved anxiogenic-like behavior that was otherwise impaired by moderate trauma in the injured WT controls (Figure 4E-F). As previously reported^28^, neither moderate CCI nor *Nlrp3* deficiency affected gross locomotor function in the OF, as shown by comparable velocity and percentage of time exploring the arena center (Figure 4 G-H). We next used NOLR task to assess short-term memory at 14-dpi. Injured *Nlrp3^-/-^* mice displayed significantly higher preference index and longer exploration time of the object in a novel location comparable to injured WT animals (Figure 4I-J). To assess long-term cognitive abilities 2 to 4 weeks after moderate CCI, we choose a more demanding behavioral paradigm. We used MWM to test sensorimotor skills, spatial- and cued learning, as well as long-term spatial memory between 18-26 dpi. During the spatial acquisition, we trained the mice for 5 days (18-22 dpi) to find a hidden platform under opaque water guided only by distal visual cues. In comparison to injured WT controls, injured *Nlrp3^-/-^* mice showed significant shorter cumulative distance to the platform (Figure 5B) and a lower percentage of time exploring the walls over time (thigmotaxis, Figure 5E). We tested the long-term spatial memory 24 hours after the last acquisition day (23 dpi) in a probe trial without platform. We observed that injured *Nlrp3^-/-^* mice crossed the virtual platform statistically similar to sham controls (Figure 5I), suggesting a trace of long-term spatial memory as observed in averaged heat maps (Figure 5H). Nevertheless, injured *Nlrp3^-/-^* mice lack precise spatial long-term memory as they show a non-significant difference in the percentage of target quadrant occupancy in comparison to another quadrant (Figure 5J).

**Figure 4.**
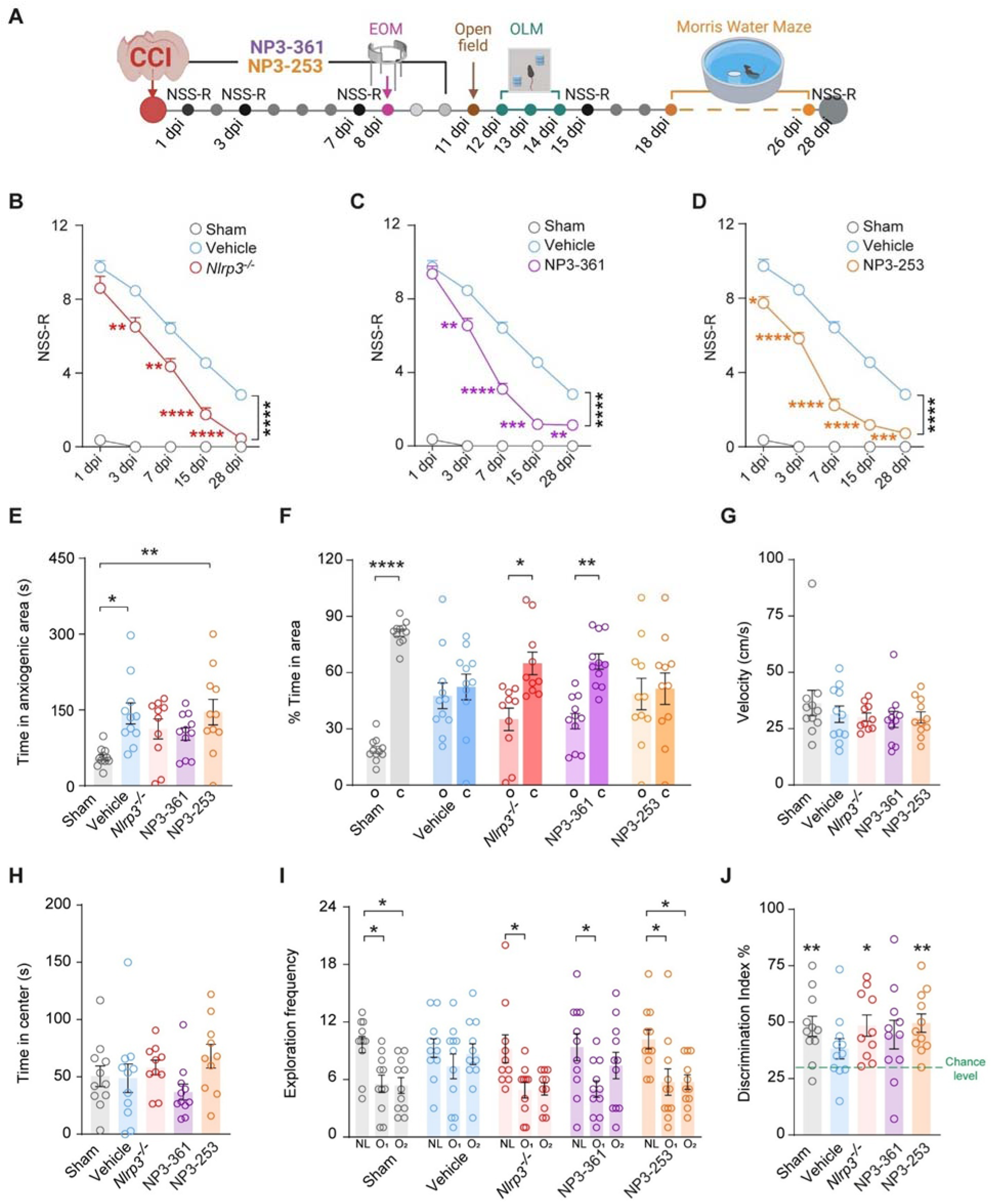
Nlrp3 gene deficiency and NLRP3 inhibition accelerate neurological recovery, and prevents cognitive impartment after CCI. (A) Schematic representation of experimental design and neurobehavioral battery. (B-D) *Nlrp3^-/-^* mice and animals treated with (C) NP3-361 or (D) NP3-253 exhibit significantly accelerated neurological recovery, as assessed by the Revised Neurobehavioral Severity Scale (NSS-R) for rodents. Two-way ANOVA with repeated measures followed by Tukey’s post hoc test. (E-F) Anxiogenic-like behavior following CCI is attenuated in *Nlrp3^-/-^* mice and in animals treated with NP3-361, as indicated by increased time spent in the anxiogenic zone of the Elevated O Maze (EOM) (E). one-way ANOVA with Tukey’s post hoc test. (F) *Nlrp3^-/-^* and NP3-361 treated animals also spent significantly more time in the closed area compared to the open areas. Paired t-test (open vs. closed areas). (I-J) Short-term spatial memory is preserved in *Nlrp3^-/-^* mice and in animals treated with NLRP3 inhibitors after severe brain injury, as shown by (I) increased exploration time of a familiar object in a novel location in the novel object location recognition test (Friedman test with Dunn’s multiple comparisons test), and (J) a higher discrimination index compared to chance level (33.33%). Wilcoxon signed-rank test. n = 10–11 mice per group. Each data point represents a biological replicate. Data are presented as mean ± SEM. Significance is indicated in red for genotype effects and in black for interaction effects. *p < 0.05, **p < 0.01, ***p < 0.001, ****p < 0.0001.

**Figure 6.**
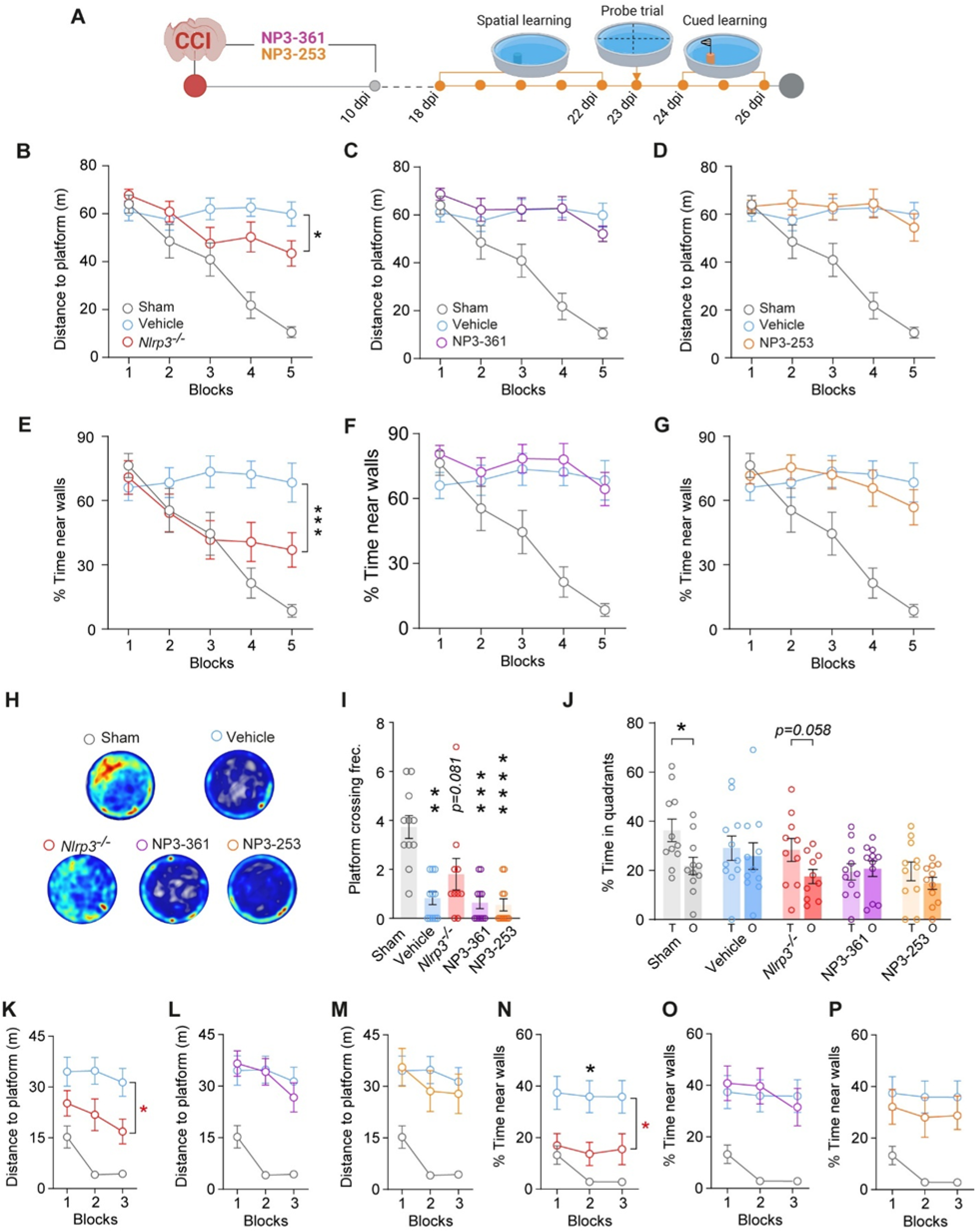
Protective effects of Nlrp3 deletion on spatial and cued learning following CCI. (A) Schematic representation of experimental design and behavioral testing on the Morris water maze paradigm. (B-D) CCI-Injured *Nlrp3^-/-^* mice display significantly shorter cumulative distances to the platform compared to CCI-Injured WT controls. Mice treated for 10 days post-injury with (C) NP3-361 or (D) NP3-253 and trained 7 days after treatment cessation exhibit cumulative distances similar to vehicle-treated injured animals. Two-way ANOVA with repeated measures followed by Tukey’s post hoc test. (E-G Injured *Nlrp3^-/-^* mice spend less time near the walls (thigmotaxis) over time compared to injured WT vehicle-treated mice. Mice treated with (C) NP3-361 or (D) NP3-253 show no significant differences relative to vehicle-treated animals. Two-way ANOVA with repeated measures and Tukey post hoc test. (H-I) Representative heat maps (H) and quantification of platform crossing frequency (I) indicate that *Nlrp3^-/-^* mice retain long-term spatial memory, as no significant differences are observed compared to sham controls. Kruskal-Wallis test with Dunn’s multiple comparisons test. (J) Impaired spatial long-term memory in injured *Nlrp3^-/-^*mice and in animals treated with NLRP3 inhibitors indicated by reduced percentage of time spent in the target quadrant versus the opposite quadrant. Wilcoxon matched-pairs signed rank test (T vs. O). (K-M) During the cued learning phase with a visible platform, injured Nlrp3-/-mice show significantly shorter cumulative distances to the platform, whereas mice treated with (L) NP3-361 or (M) NP3-253 show no significant differences relative to vehicle-treated animals. Two-way ANOVA with repeated measures followed by Tukey’s post hoc test. (N-P) injured *Nlrp3^-/-^* mice show lower percentage time near to wall (thigmotaxic) in comparison to injured WT controls, whereas mice treated with (O) NP3-361 or (P) NP3-253 show no significant differences relative to vehicle-treated animals. n = 10-11 mice per group. Individual data points in I-J represent biological replicates. All data are presented as mean ± SEM. Significance is indicated in red for genotype effects and in black for interaction effects. *p < 0.05, **p < 0.01, ***p < 0.001, ****p < 0.0001.

To test sensorimotor and visual skills, we performed an additional 3-day training with cued platform withing 24 and 26 dpi (Figure 5A and Figure 5K-P). In this phase, the mice escape from the pool by locating the platformed tagged with a striped flag. Sham WT mice learnt to locate the flagged platform with 4 trials per day. In comparison to injured WT mice, injured *Nlrp3^-/-^*mice learnt to find the platform with shorter cumulative distances (Figure 5K) and lower exploration time of the walls (Figure 5N), suggesting a preservation of sensorimotor and visual skills. Of note, naïve *Nlrp3^-/-^* and WT mice showed similar behavior in the behavioral tests conducted in the present study, consistent with previously published results^29^ and with findings from a separate, independently tested group of animals (Figure S3).

### Pharmacological treatment with oral NLRP3 inhibitor improves acute neurocognitive outcomes

Previous studies using NLRP3 modulators and intraperitoneal NLRP3 antagonists have shown potential of pharmacologically targeting of this molecule to improve peripheral and local inflammation, and neurocognitive outcome after experimental TBI. In the present study, we additionally hypothesized that acute oral treatment with a recently developed NLRP3 antagonist in the early phase after brain trauma (first 10 days) could improve long-term neurobehavioral outcomes after moderate CCI. For this purpose, we used oral NLRP3 inhibitors (NP3-253, NP3-361) with different BBB penetrance. NP3-253 possess better BBB penetration and both antagonists are administered in food pellets. Accordingly, we treated two additional groups of WT mice starting right after CCI. The treatment lasted for 10 days. We assed these groups with the same battery of behavioral tests together with the previously described injured *Nlrp3^-/-^*and vehicle-treated sham and injured groups (Figure 4A and Figure 4B). We found no evident side effects or signs of sickness behavior during the treatment with these inhibitors. Furthermore, body weight remained stable during the treatment period, however NP3-361 treated animals display lower body weights in comparison to vehicle-treated CCI control group (Figure S4 C-D). Interestingly, treated mice showed a higher intake of the food pellets with both compounds in comparison to the animals in vehicle diet (Figure S4 E-F).

In agreement with our data from injured *Nlrp3^-/-^* mice, NP3-253 and NP3-361 treated animals recovered neurological functions on NSS-R significantly faster over time (Figure 4B-C). Only NP3-361 treated and like injured *Nlrp3^-/-^* mice showed preserved anxiogenic-like behavior in the EOM expressed by the increased time in axiogenetic zone and significant longer time in closed areas (Figure 4E-F). We found a normal gross locomotor function in the OF shown by comparable velocity and percentage of time exploring the center of the arena for both inhibitors (Figure 4G-H). In the NOLR, both NP3-253 and NP3-361 treated mice were able to explore with higher exploration time the object in a novel location. Discouraging our hypothesis, we found no differences in NP3-253 and NP3-361 treated mice in comparison to vehicle treated injured controls in any of the parameters of MWM (Figure 5A-P). These results suggest a reactivation of NLRP3 and likely secondary damage and neuroinflammation. A chronic treatment with these molecules may be necessary to maintain neuroprotective effects in the long-term after trauma.

### Pharmacological inhibition of NLRP3 prevents microglial morphological response *in vivo* following moderate CCI

To determine whether pharmacological inhibition of NLRP3 replicates the effects of genetic ablation on microglial morphological responses *in vivo*, we used CX_3_CR-1*^GFP/+^*mice to visualize microglia. We then monitored CCI-induced microglial reactivity longitudinally using repeated intravital 2PLM. For this purpose, we implanted a cranial window on 6-month-old CX_3_CR-1*^GFP/+^* mice. After a three-week recovery period, we initiated treatment with NP3-361 or a vehicle diet two days before CCI induction (−2dpi, Figure 6A). To evaluate potential side effects of NP3-361 on homeostatic microglia, we performed baseline (BL) imaging one day after treatment began (−1dpi Figure 6A-B). Following baseline acquisition, we induced CCI in the hemisphere contralateral to the cranial window and conducted longitudinal imaging at 3-, 7-, and 14-dpi.

**Figure 6.**
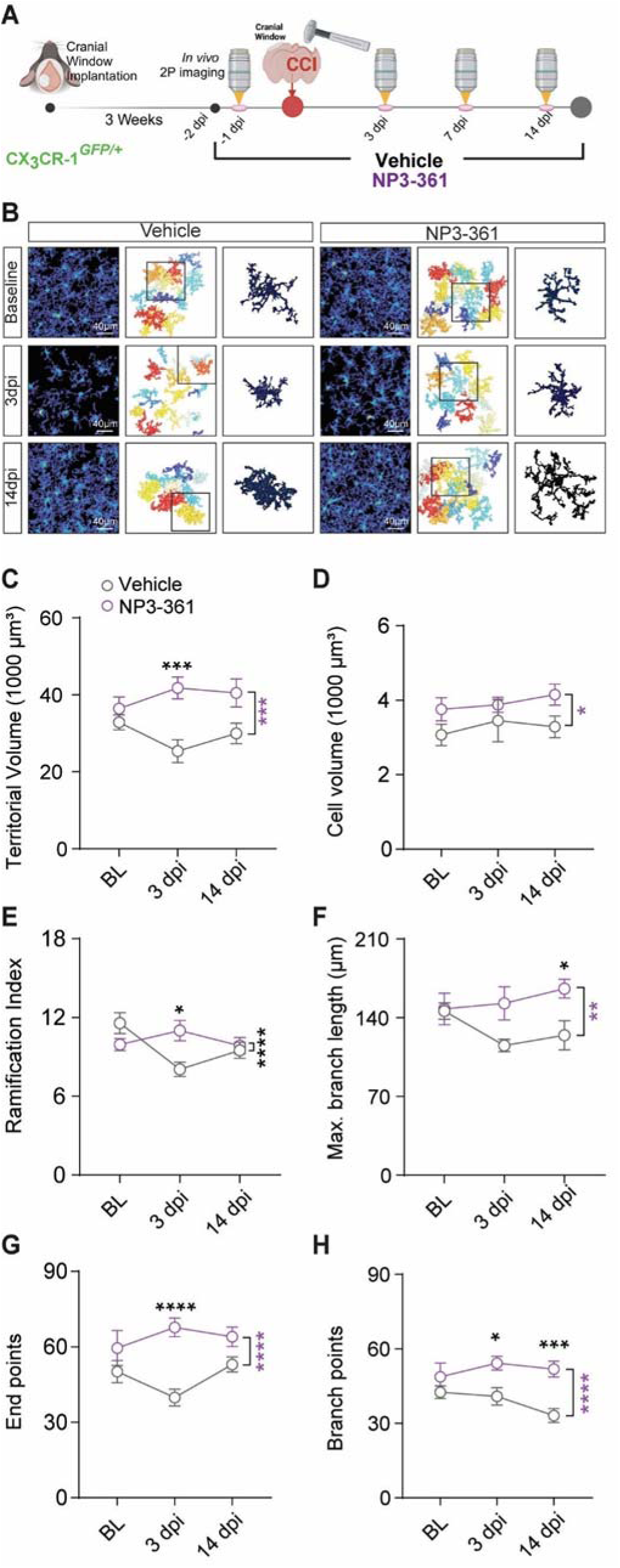
NLRP3 inhibition maintains homeostatic microglial architecture following CCI *in vivo*. (A) Schematic representation of experimental design. Cranial window surgery on *CX3CR-1^GFP/+^* 6-month-old mice, 3 weeks later treatment with NHY3619 or vehicle diet (−2dpi) was initiated. Baseline (BL) was performed one day after treatment began (−1dpi). CCI was induced in contralatel hemisphere to the cranial window and imaged longitudinally at 3, 7 and 14 dpi. (B) Representative thresholded images after repeated 2PLM imaging, 3D reconstructions, and examples of individual microglia processed and analyzed using 3DMorph^27^. (C) NP3-361 treatment prevents TBI-induced reductions in territorial volume. (D) Microglia in NP3-361 treated *CX3CR-1^GFP/+^* mice display increased somata volume across time of imaging. (E) Microglia from NP3-361 treated *CX3CR-1^GFP/+^* mice exhibit a significantly higher ramification index 3 day post-TBI. (F) Maximal branch length is significantly greater in NP3-361 treated *CX3CR-1^GFP/+^* mice at 14 days post-TBI. (G) NP3-361 treated microglia show a significantly higher number of endpoints at 3 and 14 days following CCI. (H) Higher number of branch points in NP3-361 treated microglia at 3 and 14 days post-CCI. Two-way ANOVA with repeated measures followed by Bonferroni’s post hoc test n = 16 Regions of Interest (ROI) from 4 mice per group. All data are presented as mean ± SEM. Significance is indicated in purple for treatment effect and in black for interaction effect. *p < 0.05, **p < 0.01, ***p < 0.001, ****p < 0.0001.

At baseline, NP3-361-treated animals exhibited no major morphological alterations in microglia (Figure 6C-H). Interestingly, microglia in these animals tend to display increased cell volumes and higher ramification indices compared to those in animals on a control diet (Figure 6E). Consistent with our immunohistochemical findings in *Nlrp3^-/-^* mice, acute pharmacological inhibition with NP3-361 prevented injury-induced changes in territorial volume (Figure 6D), ramification index (Figure 6E), branch length (Figure 6F), number of endpoints (Figure 6G), and branch points (Figure 6H). Notably, several of these morphological parameters remained significantly elevated even 14 days after contralateral injury, indicating preservation of a homeostatic microglial state under NP3-361 treatment. Our longitudinal imaging approach thus demonstrates *in vivo* that NLRP3 inhibition not only prevents cognitive impairment but also suppresses microglial reactivity following injury, mirroring the effects observed with genetic ablation of NLRP3. These findings align with recent evidence showing that pharmacological NLRP3 inhibition can directly modulate microglial morphology, shifting cells from a hyperactivated ameboid state to a more homeostatic, ramified phenotype.

## DISCUSSION

We confirmed that NLRP3 and related inflammasome proteins are consistently upregulated during the initial phases after TBI (Figure 1C-L). This upregulation promotes ASC aggregation (Figure 2 A-E) and is essential for both acute and long-term microglial morphological changes (Figure 3 A-G and Figure 6 B-H), as well as for the neurobehavioral impairment following brain injury. Notably, we demonstrated that *Nlrp3^-/-^* mice show accelerated neurological recovery and improved neurocognitive outcomes, not only in acute but also late phases of TBI. Additionally, we tested newly developed oral NLRP3 inhibitors and found that, at least during the period of administration, mice subjected to experimental TBI exhibited improved neurocognitive function (Figure 4 and Figure 5). Furthermore, using repeated longitudinal intravital imaging *in vivo*, we confirmed that treatment with an NLRP3 inhibitor preserved microglial homeostatic features in TBI-injured mice (Figure 6 B-H).

Previous studies have used a weight-drop model to induce moderate to severe CHI in *Nlrp3^-/-^*mice^17,18,30^, but the findings have been inconsistent. While some researchers observed reduced IL-1β and CASP1 activity post-injury in these mice, others did not detect significant changes in gene expression related to these proteins^17,18,30^. Additionally, outcomes such as brain water content^30^, short-term novel object recognition memory^17^, blood-brain barrier integrity^18^, and neurological scores varied between studies, possibly due to differences in the experimental protocols employed. In the present study, we observed a significant impairment in the upregulation of the effector protein CASP1 and the mediators GSDMD, IL-1β, and IL-18 in *Nlrp3^-/-^* mice (Figure 1). Despite these significant genotype-dependent differences, a complete suppression was not observed, likely due to compensatory activation of other types of inflammasomes, such as AIM2^18,31^. This observation may indicate reduced pyroptosis in *Nlrp3^-/-^ mice*, which we confirmed in our histological analysis by detecting ASC aggregates (Figure 2). We also observed smaller lesion sizes in *Nlrp3^-/-^* mice after CCI, which persisted up to 30 days post-injury (Figure 1A-B). Although our study employed a different model, this finding aligns with the reduced histopathological abnormalities reported before^17^ and the results obtained following prolonged intraperitoneal treatment with the NLRP3 inhibitor MCC950^22^. An anti-apoptotic effect has been proposed to explain these phenomena, based on differential expressions of apoptotic markers such as cleaved caspase-3, Annexin V, or BAX in injured *Nlrp3^-/-^* or treated mice. However, we detect no significant difference in apoptosis using the TUNEL assay in *Nlrp3^-/-^* mice at 3 days post-injury (Figure S1). Although crosstalk between apoptotic and inflammasome pathways has been previously described^32^, in our model, the absence of the *Nlrp3* gene does not appear to significantly affect the occurrence of this type of cell death after CCI. We propose that the reduced lesion size in *Nlrp3^-/-^* mice may be primarily explained by modulation of pyroptosis pathways, as we observed lower ASC aggregation both within and outside microglia (Figure 2).

Microglia are key players in the brain’s neuroinflammatory response. Whether they exert protective or detrimental effects during secondary damage following traumatic injury remains controversial. A classical and reliable indicator of microglial activity is their morphology. Since NLRP3 is predominantly expressed by microglia^33^, and *Nlrp3^-/-^* mice exhibit smaller lesions and improved neurocognitive outcomes, we analyzed the detailed morphology of *Nlrp3^-/-^* microglia at various time points following CCI in the perilesional tissue (Figure 3). We found that *Nlrp3^-/-^* microglia displayed significantly fewer features characteristic of activated ameboid or rod-like morphologies, as previously described in a CHI model. This observation aligns with our earlier findings on microglial morphology following systemic inflammation in the APP/PS1 model^34^. We demonstrated that *Nlrp3^-/-^* microglia maintain a resilient architecture in response to inflammatory stimuli such as LPS or direct mechanical trauma. This resilient morphology may be associated with an enhanced microglia-specific genetic program, as described by previous reports, including overexpression of genes such as *Clec7a*, *Cd11b*^18^, and *C1qa* (Figure S2R). Future genomic and proteomic analyses of isolated microglia after TBI may elucidate the specific mechanisms underlying this resilient phenotype.

After finding that *Nlrp3^-/-^* mice exhibited smaller lesions, resilient microglial architecture and reduced ASC aggregation, we decided to conduct a behavioral characterization of *Nlrp3^-/-^* mice following TBI (Figure 4 and Figure 5). We performed a battery of standardized behavioral assessments over long term. It was previously reported that *Nlrp3^-/-^* mice maintain preserved short-term recognition and working memory at later stages (4 and 8 weeks) after CCI^35^. We not only replicated these observations but also assessed more complex paradigms, such as the Morris Water Maze (MWM), in which CCI-injured *Nlrp3^-/-^*mice demonstrated some preserved cognitive abilities. We further evaluated the potential of oral NLRP3 pharmacological inhibitors to replicate the behavioral phenotype observed in *Nlrp3^-/-^* mice following TBI. We hypothesized that administering these inhibitors during the first 10 days post-TBI could ultimately preserve cognitive function at later stages. Notably, both tested oral inhibitors (NP3-253 and NP3-361) exhibited protective effects during the treatment period comparable to those seen with *Nlrp3* genetic deficiency. Although NP3-361 presumably possess poor BBB penetration, its beneficial effects may be attributed to the pronounced BBB disruption that accompanies TBI or to the involvement of peripheral effects. Additionally, we found that this protective effect was no longer evident eight days after cessation of inhibitor treatment, suggesting that NLRP3-mediated neuroinflammation was likely reactivated. This indicates that chronic or continuous inhibition may be necessary to sustain protection after TBI, consistent with previous findings using prolonged intraperitoneal MCC950 administration^22^. Similar protective effects with chronic NLRP3 inhibition were recently observed in an *in vitro* model of AD-related neuroinflammation^36^. To assess the longitudinal, *in vivo* effects of pharmacological NLRP3 inhibition on microglial morphology as a cellular proxy of neuroinflammation, we conducted repeated intravital 2PLM in NP3-361–treated animals, both before and up to 14 days following CCI (Figure 6). Our results confirmed the protective effects on microglial reactivity and the preservation of homeostatic morphological features, consistent with observations in *Nlrp3^-/-^*mice. Comparable protective effects on microglial morphology have also been reported with chronic NLRP3 inhibition using MCC950 in models of chronic unpredictable mild stress^37^ and subarachnoid hemorrhage^38^, as well as with the BBB-penetrant NLRP3 inhibitor VEN-02XX in an AD model^39^.

Overall, our results underscore the critical role of NLRP3 in the neuroimmunological response to TBI, particularly in mediating changes in microglial architecture, promoting ASC aggregation and processing of neuroinflammatory mediators. These findings highlight NLRP3 as a promising therapeutic target and lay the groundwork for the development of pharmacological interventions aimed at modulating the NLRP3 inflammasome in TBI. Such interventions could be beneficial in both the acute and chronic management of patients who have experienced TBI or are at risk of developing TBI-related sequelae. Furthermore, our data support that targeting NLRP3 in TBI attenuates neuroinflammatory cascades, reduce secondary damage and improve neurological outcomes.

### Limitations of the study

Throughout this study, we primarily attribute the protective effects observed following *Nlrp3* deletion or pharmacological inhibition of NLRP3 to local neuroinflammatory mechanisms, particularly microglial inflammasome activation. However, we cannot fully exclude the contribution of NLRP3 signaling in other CNS cells, or peripheral immune cells, nor can we disregard the involvement of other organs in the systemic response to TBI. For instance, recent evidence indicates that *nlrp3* ablation in astrocytes can contribute to behavioral phenotypes observed after mild TBI, suggesting a broader cellular involvement in the pathophysiology of TBI than previously appreciated^40^. Given that TBI induces a systemic inflammatory response and involves multiple organ systems, our study investigates both global *Nlrp3* deletion and systemic administration of NLRP3 inhibitors, as systemic inhibition may offer a promising therapeutic strategy to limit the widespread, multi-organ inflammation associated with TBI. However, further research is required to clarify the distinct roles of NLRP3 in various CNS and peripheral cell populations, and to better understand the mechanisms that differentiate systemic from local effects. Future studies should also thoroughly assess the safety, optimal dosing, and therapeutic efficacy of NLRP3 inhibitors in patients after TBI in controlled clinical trials.

## RESOURCE AVAILABILITY

### Lead contact

Further information and requests for resources and reagents should be directed to and will be fulfilled by the lead contact, Michael T. Heneka.

### Materials availability

This study did not generate new unique reagents.

### Data and code availability

- All data needed to support the conclusions are presented in the paper or the supplemental information. Additional data related to this paper may be requested from the authors.
- This study does not report any custom code.
- Any additional information required to reanalyze the data reported in this work paper is available from the lead contact upon request.

## ACKNOWLEDGMENTS

The NLRP3-specific inhibitor ‘‘NP3-361’’ and ‘‘NP3-253’’ were gifts from Novartis. We thank I. Rácz and C. Ising for help with obtaining approval by the local ethical committee for the animal experiments. Sean-Patrick Hermann for assistance with editing the final draft; Paula Martorell, and Nadia Villacampa for valuable discussions; and the DZNE Light Microscopy Facility for support with imaging acquisition. S.C-G., F.D.W., F.M; E.L., M.T.H. are members of the excellence cluster ImmunoSensation funded by DFG Germany’s Excellence Strategy–EXC2151–390873048. S.C.-G. is supported by Alzheimer Forschung Initiative e.V (Grant 21060), the Hertie Network of Excellence in Clinical Neuroscience (Grant 2021-1A-12), the BONFOR-Forschungskommission der Medizinischen Fakultät Bonn and by the Deutsche Forschungsgemeinschaft (DFG, German Research Foundation) under Germany’s Excellence Strategy– EXC2151–390873048, and the Neuro-aCSis Bonn Neuroscience Clinician Scientist Program.

## AUTHOR CONTRIBUTIONS

Conceptualization, S.C-G., L.E. and M.T.H; methodology, S.C-G., A.V-S., S.S. and F.M.; Investigation, S.C-G., A.V-S., S.S., J.I.M-M., I.K., F.D.W.; writing—original draft, S.C-G and M.T.H.; writing—review & editing, S.C-G and M.T.H.; funding acquisition, S.C-G., V.S. and M.T.H..; resources, S.C-G, E.L., V.S. and M.T.H.; supervision, S.C-G and M.T.H.

## DECLARATION OF INTERESTS

M.T.H. is a scientific advisory board member at Alector the Dementia Discovery Fund, and Muna Therapeutics and has received honoraria for oral presentations from Pfizer, Novartis, Roche, Abbvie, and Biogen. E.L. is a co-founder and adviser at IFM Therapeutics, Dioscure Therapeutics, Stealth’’ Biotech, and Odyssey Therapeutics.

## DECLARATION OF GENERATIVE AI AND AI-ASSISTED TECHNOLOGIES

The authors used perplexity.ai to check for grammar and style.

## STAR METHODS

Detailed methods are provided in the online version of this paper and include the following:

- KEY RESOURCES TABLE
- EXPERIMENTAL MODEL AND STUDY PARTICIPANT DETAILS
- METHOD DETAILS

- Controlled cortical impact (CCI)
- Experimental design and drug administration
- Neurobehavioral evaluation
- Tissue preparation
- Histology and immunohistochemistry
- TUNEL staining
- Quantitative analysis of ASC aggregates and microglia morphology
- Protein extraction
- Protein Extraction and Immunoblot analysis
- ELISA pro-inflammatory response quantification
- RNA preparation and RT-PCR
- Cranial window surgery
- Intravital two-photon laser scanning microscopy
- QUANTIFICATION AND STATISTICAL ANALYSIS

- Statistics and Reproducibility

## STAR METHODS

## KEY RESOURCES TABLE

| REAGENT OR RESOURCE | SOURCE | IDENTIFIER |
| --- | --- | --- |
| **Antibodies** |  |  |
| Anti Iba1, Rabbit | FUJIFILM Wako Pure Chemical Corporation | Cat. No. 01919741<br>RRID: AB_839504 |
| ASC/TMS1 (D2W8U) Rabbit mAb | Cell Signaling | Cat. No. 67824<br>RRID: AB_2799736 |
| Cleaved Gasdermin D (Asp276) (E3E3P) Rabbit mAb | Cell Signaling | Cat. No. 10137<br>RRID: AB_2923068 |
| NLRP3 (D4D8T) | Cell Signaling | Cat. No. 15101,<br>RRID: AB_2722591 |
| Rat Caspase-1 Antibody, clone 4B4.2.1 | Gift from Genentech | N/A |
| Anti-Caspase-1 (p20) (human) mAb (Bally-1) | AdipoGen | Cat. No. AG-20B-0048,<br>RRID: B_2490257 |
| Anti-Caspase-1 (p20) (mouse) mAb (Casper-1) | AdipoGen | Cat No. AG-20B-0042; RRID: AB_2490248 |
| Alexa Fluor 488-AffiniPure Donkey Anti-Goat IgG | Jackson ImmunoResearch | Cat No. 705-545-147,<br>RRID: AB_2336933 |
| Alexa Fluor 647-AffiniPure Donkey Anti-Goat IgG | Jackson ImmunoResearch | Cat No. 706-605-003,<br>RRID: AB_2340436 |
| Alexa Fluor 488-AffiniPure Donkey Anti-Mouse IgG | Jackson ImmunoResearch | Cat No. 715-545-150,<br>RRID: AB_2340846 |
| Donkey Anti-Mouse IgG H&L (Alexa Fluor® 555) | Abcam | Cat No. ab150110,<br>RRID: AB_2783637 |
| Donkey anti-rabbit Alexa Fluor 555 | Invitrogen | Cat No. A-31572,<br>RRID: AB_162543 |
| β-Actin Antibody | Cell Signaling | Cat No. 4967, RRID: AB_330288 |
| IRDye 680LT Goat anti-Mouse IgG | LI-COR Biosciences | Cat No. 926-68020,<br>RRID: AB_10706161 |
| IRDye 800CW Goat anti-Mouse IgG | LI-COR Biosciences | Cat No. 925-32210,<br>RRID: AB_2687825 |
| IRDye 680RD Goat anti-Rat IgG | LI-COR Biosciences | Cat No. 926-68076,<br>RRID: AB_10956590 |
| IRDye 800CW Goat anti-Rabbit IgG | LI-COR Biosciences | Cat No. 926-32211,<br>RRID: AB_621843 |
| **Chemicals, peptides, and recombinant proteins** |  |  |
| Cresyl violet solution | Sigma C-1791 | Cat No. C-1791 |
| Entellan™ | Merck | Cat No.107960 |
| DAPI | Thermo Fischer Scientific | Cat No. D1306 |
| Fetal Bovine Serum | Life Technologies | Cat No. 10270106 |
| NP3-253 | Gift from Novartis | N/A |
| NP3-361 | Gift from Novartis | N/A |
| NuPAGE 4–12% Bis-Tris gel | Invitrogen | Cat No. NP0323BOX |
| Paraformaldehyde | Sigma-Aldrich | Cat No. P6148 |
| Qiazol lysis reagent | Qiagen | Cat No. 79306 |
| TaqMan universal PCR master mix | Thermo Fisher Scientific | Cat No. 4304437 |
| Immu-Mount™ | Epredia | Cat No. 9990402 |
| Critical commercial assays |  |  |
| Pierce BCA Protein Assay kit | Thermo Fisher Scientific | Cat No.23225 |
| V-PLEX Pro-inflammatory Panel 1 Mouse Kit | Meso Scale Discovery | Cat No. K15048D-1 V-PLEX |
| Click-iT™ Plus TUNEL Assay | Thermo Fisher Scientific | Cat. No. C10617, C10618, C10619 |
| Experimental models: Organisms/strains |  |  |
| Mouse: C57BL/6N | Charles River | RRID: ISMR_CRL:027 |
| Mouse: *CX3CR-1<sup>GFP/+</sup>* | The Jackson Laboratory | RRID: IMSR_JAX:005582 |
| Mouse: *Nlrp3<sup>-/-</sup>* | Millennium Pharmaceuticals<sup>41</sup> | N/A |
| TaqMan® Gene Expression Assays |  |  |
| *Gapdh* | Thermo Fisher Scientific | Mn99999915_g1 |
| *C3* | Thermo Fisher Scientific | Mm00437858_m1 |
| *C1aq* | Thermo Fisher Scientific | Mm00432142_m1 |
| *Il-18* | Thermo Fisher Scientific | Mm00434226_m1 |
| Software and algorithms |  |  |
| 3DMorph | York et. al<sup>27</sup> | https://github.com/ElisaYork/3DMorph |
| Ethovision XT15 | Noldus | https://www.noldus.com/ethovision-xt |
| Fiji ImageJ | GNU General Public License | https://imagej.net/software/fiji/ |
| ImageStudio software | LI-COR Biosciences | www.licor.com/bio/products/software/image_studio_lite |
| Imaris v 9.5 | Oxford Instruments | https://imaris.oxinst.com/ |
| MATLAB | Mathwork | https://www.mathworks.com/products/matlab.html |
| NIS-Elements | Nikon | https://www.microscope.healthcare.nikon.com/products/software/nis-elements |
| Prism 9.5.0 | GraphPad | https://www.graphpad.com/scientific-software/prism/ |
| R Studio v.4.5 | Posit software | https://rstudio.com/ |
| ZEN 2012 Blue Edition | Zeiss | https://www.zeiss.com/microscopy/en/products/software/zeiss-zen.html |
| Other |  |  |
| Nikon A1R MP + multiphoton | Nikon | Equipment |
| Leica SM2010R Sliding Microtome | Leica | Equipment |
| NanoDrop | Thermo Fisher Scientific | Equipment |
| LI-COR ODYSSEY CLx Imaging System | LI-COR Biosciences | Equipment |
| QuickPlex SQ 120 plate reader | Meso Scale Discovery | Equipment |
| StepOnePlus Real-Time PCR System | Applied Biosystems | Equipment |
| Zeiss Axio Scan.Z1 | Zeiss | Equipment |
| Zeiss LSM 800 confocal microscope | Zeiss | Equipment |

## EXPERIMENTAL MODEL AND STUDY PARTICIPANT DETAILS

Male and female *Nlrp3^-/-^* mice (6–7 months old) and wild-type (WT) littermates on a mixed C57BL/6NJ genetic background were used1. *Nlrp3^-/-^* mice were obtained from Millennium Pharmaceuticals and previously described^41^. Mice were housed in groups of 2–4 animals per cage in animal facilities at the the University Hospital Bonn under a 12-h light/dark cycle, with *ad libitum* access to food and water, and under specific pathogen-free conditions. Behavioral experiments were conducted during the dark phase (09:00–19:00 h). Animal procedures followed the ARRIVE guidelines and were approved by the German Animal Welfare Authorities of the North Rhine-Westphalia government (approval numbers 81-02.04.2019.A026 and 81-02.04.2021.A197)

## METHOD DETAILS

### Controlled cortical impact (CCI)

CCI was employed to model moderate TBI. Surgical procedures were conducted under inhaled anesthesia. Mice were initially anesthetized with 5% isoflurane delivered in 100% oxygen (flow rate: 0.5–1 L/min) for induction and subsequently maintained at 1.5–2% isoflurane throughout the surgical procedure. Animals were positioned in a stereotaxic frame (Stoelting, Dublin, Ireland). After making a midline scalp incision, a circular craniotomy (4–5 mm in diameter) was performed over the left parietal cortex using a motorized 0.4 mm dental drill (Schick, Schemmerhofen, Germany). CCI was induced using an electromagnetic impactor (Impact One Stereotaxic Impactor, Leica Biosystems, Germany) with a 3 mm diameter metal tip. The impactor was programmed with a velocity of 5 m/s, a dwell time of 200 ms and a depth of 1.5 mm. The lesion epicenter was located 2 mm lateral to the midline and 2 mm posterior to bregma. Sham-operated mice underwent identical anesthesia and surgical preparation, but without undergoing craniotomy or cortical injury. Postoperative analgesia was provided by subcutaneous administration of carprofen (5 mg/kg, once daily) and by supplementing the drinking water with tramadol (0.1 mg/mL) for three days following surgery.

### Experimental Design and Drug Administration

A total of 44 WT mice were randomized into four groups (n = 11 per group): Sham-operated controls, CCI-injured WT mice receiving standard diet (Cehicle), CCI-injured WT mice treated with NP3-361 (50 mg/kg), and CCI-injured WT mice treated with NP3-253 (180 mg/kg). Both NP3-361 and NP3-253 were formulated in the standard diet. An additional group of 10 *Nlrp3^-/-^* mice underwent CCI surgery and was tested in parallel. Drug administration began immediately after CCI surgery, with inhibitors provided in food pellets. The vehicle group received only the standard diet.

### Neurobehavioral evaluation

#### Revised Neurobehavioral Severity Scale (NSS-R)

The NSS-R is a sequence of neurological evaluations designed to assess sensory and motor functions, including general movement, including reflex suppression, and postural adjustments. Scores on the NSS-R range from 0 (normal) to 20 (maximal deficit)^42^. Assessments were performed prior to surgery and at 1, 3, 7, 14, and 28 days following CCI or sham surgery.

#### Elevated O maze (EOM)

Anxiety-like behavior and risk assessment were assessed using an EOM, consisting of an annular runway (diameter: 46 cm; width: 5.5 cm) elevated 40 cm above the ground, with two wall-protected sectors alternating with two open sectors.

#### Open Field

Spontaneous exploration and locomotor activity were assessed in an open field arena (50 cm × 50 cm × 50 cm) built with opaque gray acrylic. For each test, mice were video recorded for 15 minutes under indirect dim illumination (40 lux).

#### Novel object location recognition test (NOLR)

The NOLR test was conducted in the open-field box as previously described. The floor of the arena was covered with a 1-cm layer of sawdust that had been previously used and saturated with the animals’ odor. During the habituation period, animals were allowed to freely explore the arena for 5 minutes per day over 2 consecutive days. Three identical objects (Lego pieces of different colors, approximately 2 × 4 cm) were placed in fixed locations within the arena during habituation.On the test day, animals were first allowed to explore the three identical objects for 6 minutes. The time spent investigating each object was recorded using an automated tracking system (Noldus Information Technology, Wageningen, Netherlands). One hour later, one of the objects was relocated to a novel position within the arena, while the other two remained in their original locations. Animals were then returned to the arena and allowed to explore for 5 minutes. The time spent investigating each object, beginning with the first contact, was recorded. The preference ratio for the moved object was calculated for the first 3 minutes using the following formula: Discrimination Index (%) = [Time spent exploring the object in the novel location / Total time spent exploring all objects] × 100.

#### Morris water maze (MWM)

Spatial learning and memory were assessed 18–28 days after CCI using the MWM. A circular pool (1 m in diameter) was filled with water rendered opaque by the addition of non-toxic white dye and maintained at 21–23 °C. The pool was dimly illuminated (∼40 lx) and enclosed by a white curtain to reduce external visual distractions. Distinct, asymmetrically positioned distal cues were attached to the pool walls to provide spatial reference points. The pool was virtually divided into four quadrants, with a hidden platform (15 cm diameter) submerged 1.5 cm below the water surface in the middle of one of the quadrants. Mice were trained over five consecutive days to locate the hidden platform using the distal cues. Each day consisted in four trials per day. In each trial, mice were introduced into the water from pseudo-randomized starting positions to discourage reliance on non-spatial strategies. Each mouse was given 60 seconds to find the platform; in case of not reaching it, the mouse was gently guided to it. After reaching the platform, mice remained there for 15 seconds before being returned to their home cage upon completion of all four trials. Twenty-four hours after the final training session, a spatial probe trial was performed. During this trial, the platform was removed, and each mouse was placed in the quadrant opposite the previous platform location, facing the wall at the midpoint of the quadrant. Mice were allowed to swim freely for 60 seconds, and their search patterns were recorded and analyzed.

Subsequently, a cued learning phase was conducted to assess visual and motor abilities. In this phase, the platform was made visible by marking it with a flag and was repositioned to a different quadrant, while all distal cues were removed. Mice underwent three trials per day for three consecutive days to locate the flagged platform. All behavioral sessions were video-recorded and analyzed using EthoVision XT (Noldus Information Technology, Wageningen, Netherlands).

### Tissue preparation

For organ harvesting, mice were anesthetized using a combination of ketamine (100 mg/kg body weight) and xylazine (20 mg/kg body weight). Following anesthesia, blood was collected from the right ventricle. The mice were then subjected to transcardial perfusion with approximately 30 mL of cold PBS. Subsequently, brains were carefully dissected. For histological assessment, the brain samples were immersion-fixed in 4% paraformaldehyde for 24 hours at 4°C. After fixation, the brains were rinsed three times with cold PBS and then stored in PBS containing sodium azide. For biochemical analysis, perilesional brain tissue was promptly dissected, rapidly frozen in liquid nitrogen, and stored at −80°C.

### Histology and immunohistochemistry

Coronal brain sections (40 µm) were prepared using a Leica VT1000S vibratome and processed as free-floating sections for subsequent staining. For Nissl staining, sections were mounted onto glass microscope slides (Thermo Scientific, Waltham, USA) and allowed to dry overnight. To remove lipids, the tissue was immersed overnight in a 1:1 mixture of absolute ethanol and chloroform. The following day, slides were rehydrated by sequential immersion in 100% ethanol, followed by 95% (v/v) ethanol. Sections were then stained in 0.1% (w/v) cresyl violet solution (C-1791, Merck, Darmstadt, Germany) for five minutes, rinsed with distilled water, and differentiated in 95% (v/v) ethanol for up to 30 minutes, with staining intensity monitored every five minutes. Once the desired staining intensity was achieved, slides were dehydrated through two incubations in 100% ethanol and two in xylene. Finally, sections were air-dried and cover slipped using Entellan mounting medium (Merck, Darmstadt, Germany). Whole-section images were acquired using a Zeiss Axio Scan.Z1 microscope equipped with a 20× objective.

For immunohistochemical analysis, sections were washed three times for five minutes each in PBS. They were then incubated in citrate buffer at 95°C for five minutes to perform antigen retrieval. After cooling to room temperature, sections were washed in PBS containing 0.5% Triton X-100 (PBS-T). Non-specific binding was blocked by incubating sections in 1% bovine serum albumin (BSA) in PBS-T for one hour. Sections were then incubated overnight at 4°C with the appropriate primary antibodies. On the following day, sections were washed three times for five minutes each in PBS-T, incubated with secondary antibody conjugates (usually in 1:500 dilution) for 60 minutes at room temperature, and washed three times for five minutes each in PBS. Finally, sections were mounted using ProLong Gold Antifade Mountant with DAPI (Thermo Scientific, Waltham, USA).

### TUNEL staining

To assess apoptotic cell presence brain sections were stained using the Click-iT™ Plus TUNEL Assay (Cat. No. C10617, C10618, C10619; Invitrogen) according to the manufacturer’s instructions. Sections were mounted with ImmuMount containing 1[mg/ml DAPI (Thermo Fisher Scientific) and coverslipped. Whole-section images were acquired using a Zeiss Axio Scan.Z1 equipped with a 20× objective.

### Quantitative analysis of ASC aggregates and microglia morphology

Images of whole brain sections High-magnification imaging were acquired using a Zeiss LSM700 confocal microscope with a 40× oil immersion objective. Image processing was performed using ImageJ (Version 2.0.0-rc-67/1.52c) and Imaris (Version 9.5). Quantification of ASC aggregates was carried out in Imaris (Version 9.5) following the manufacturer’s instructions. Three-dimensional analysis of microglial morphology and complexity was conducted using 3DMorph, an open-source MATLAB-based script, according to previously published protocols and programmer instructions^27^.

### Protein Extraction and Immunoblot analysis

For protein extraction, tissue samples were thawed and homogenized in PBS containing 1 mM EDTA, 1 mM EGTA, and a protease inhibitor cocktail (Thermo Scientific, Waltham, USA). The resulting homogenates were then subjected to further extraction using RIPA buffer, which was composed of 25 mM Tris–HCl (pH 7.5), 150 mM NaCl, 1% Nonidet P-40, 0.5% sodium deoxycholate, and 0.1% sodium dodecyl sulfate (SDS). Following extraction, samples were centrifuged at 20,000 × g for 30 minutes at 4°C. Protein samples were separated by electrophoresis on NuPAGE gels (Thermo Scientific, Waltham, USA) and subsequently transferred onto nitrocellulose membranes (Hercules, California, USA). The membranes were incubated with the following primary antibodies: anti-Caspase-1 (1:1000 dilution, Genentech, USA), anti-ASC (clone D2W8U, 1:1000, Cell Signaling, Germany), anti-NLRP3 (clone D4D8T, 1:500, Cell Signaling, Germany), and anti-Gasdermin D (clone E3E3P, 1:1000, Cell Signaling, Germany). After incubation with the appropriate secondary antibodies, immunoreactive bands were detected using the Odyssey CLx imaging system (LI-COR, Germany) and quantified using ImageJ software (NIH, USA).

### ELISA pro-inflammatory response quantification

Proinflammatory Panel 1 (Mouse) assay kits (Meso Scale Discovery, Rockville, MD, USA) were used to quantify ten cytokines: IFN-γ, IL-1β, IL-2, IL-4, IL-5, IL-6, KC/GRO, IL-10, IL-12p70, and TNF-α. All procedures were conducted following the manufacturer’s protocol. Briefly, 50 µL of diluted sample, calibrator, or control was added to each well. The plates were sealed with adhesive film and incubated at room temperature with gentle shaking for 2 hours. Following incubation, plates were washed three times, and detection antibodies were added. Plates were resealed and shaken at room temperature for an additional 2 hours. After a second wash, read buffer was applied, and signal detection was performed using a Sector Imager 2400 plate reader (Meso Scale Discovery, Rockville, MD, USA).

### RNA preparation and RT-PCR

RNA concentration and purity were assessed using a NanoDrop 1000 spectrophotometer (Thermo Scientific, Waltham, USA). For each sample, 3 µg of total RNA was reverse transcribed into complementary DNA (cDNA) with the SuperScript III Reverse Transcriptase kit (Thermo Scientific, Waltham, USA). Real-time quantitative PCR was conducted on a StepOnePlus Real-Time PCR System (Applied Biosystems, Waltham, Massachusetts, USA) utilizing TaqMan Gene Expression Assays (Thermo Scientific, Waltham, USA). Each 20 µl reaction included 10 ng of cDNA, 10 µl of 2× Gene Expression Master Mix, and 1 µl of the specific TaqMan Gene Expression Assay. The following assays were employed: Gapdh (Assay ID: Mn99999915_g1), C3 (Assay ID: Mm00437858_m1), C1qa (Assay ID: Mm00432142_m1), and Il18 (Assay ID: Mm00434226_m1). GAPDH was used as an internal control. All samples were analyzed in triplicate, and relative gene expression levels were normalized to GAPDH.

### Cranial window surgery

Mice were anesthetized with 5% isoflurane in 100% oxygen (flow rate: 0.5–1 L/min) for induction and maintained at 1.5–2% isoflurane during surgery. The animals were fixed in a stereotaxic frame (Stoelting, Dublin, Ireland). A midline scalp incision was made, followed by a circular craniotomy (approximately 5 mm in diameter) over the left parietal cortex using a motorized 0.4 mm dental drill (Schick, Schemmerhofen, Germany). A 5 mm coverslip was attached to the skull with cyanoacrylate glue (UHU, Brühl, Germany). A customized titanium ring was then fixed to the skull using dental cement (Heraeus Kulzer, Hanau, Germany). After a 3-week recovery period, mice were imaged.

### Intravital two-photon laser scanning microscopy

A Nikon A1R MP microscope, equipped with a titanium–sapphire laser (Chameleon Ultra, Coherent, Santa Clara, CA), was used to conduct intravital imaging of the mouse brain. The same group of mice underwent imaging at four distinct time points: baseline (prior to injury), and at 3, 7, and 14 days following CCI. Throughout all imaging procedures, the laser power remained below 30 mW. No animal mortality occurred during or after imaging sessions or as a result of CCI. Three-dimensional z-stacks were acquired, spanning a depth of 50 μm with a 1 μm step size between optical planes, using a Nikon 25× objective lens (1.1 NA, CFI75 Apochromat 25XC W 1300). The three-dimensional morphology and complexity of microglia were automatically quantified using the 3DMorph pipeline^27^.

## QUANTIFICATION AND STATISTICAL ANALYSIS

### Statistics and Reproducibility

All data are presented as mean ± SEM. Statistical analyses were conducted using GraphPad Prism version 9.5 or higher (GraphPad Software, Boston, United States) for data visualization and RStudio for advanced statistical analyses, including ANOVA on rank-transformed data. Prior to hypothesis testing, data distribution was assessed for normality using the D’Agostino-Pearson omnibus test. Comparisons between two groups were performed using a two-tailed unpaired t-test for normally distributed data, while the Mann-Whitney U test was applied to non-normally distributed data. For analyses of more than two groups, either one-way or two-way ANOVA was utilized. In experiments where repeated measurements were collected from the same subjects over conditions or time, repeated measures two-way ANOVA was specifically employed to account for within-subject variability. Tukey’s or Bonferroni post hoc tests were conducted as appropriate following significant ANOVA results. When data failed to meet assumptions of normality or homogeneity of variances, nonparametric alternatives were used. For ranked or ordinal data—or when normality assumptions were violated—ANOVA on ranks was performed in R by replacing original values with their rank order and applying standard ANOVA procedures to the transformed data. In factorial designs where interaction effects were of interest, the aligned rank transform (ART) ANOVA was applied to robustly test both main and interaction effects in non-normally distributed data. Alternatively, for one-way designs, the Kruskal-Wallis test with Dunn’s post hoc test was used when appropriate. p < 0.05 was considered statistically significant.

## SUPPLEMENTAL INFORMATION

**Figure S1.**
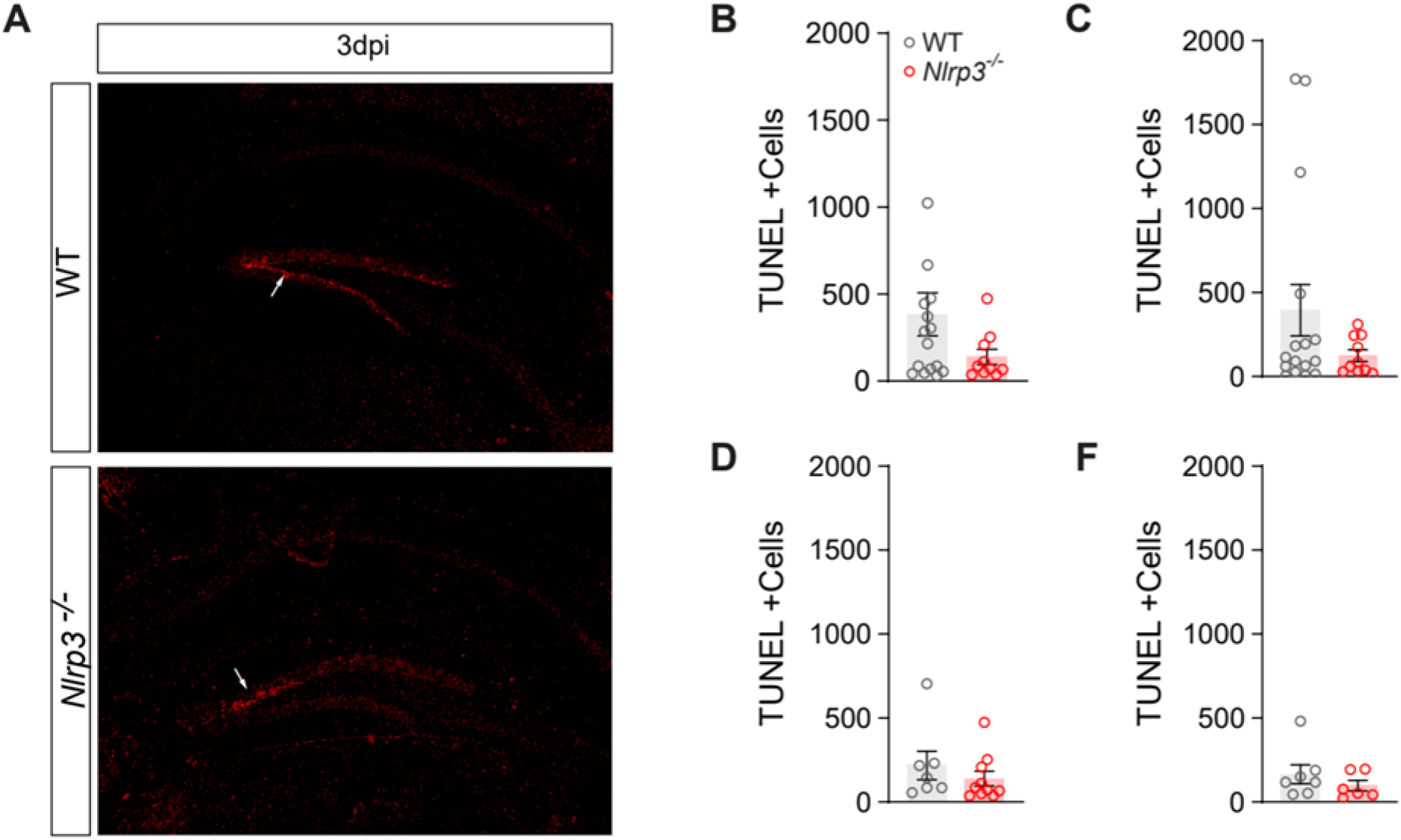
WT and *Nlrp3^-/-^* mice display similar apoptosis levels following CCI. (A) Representative images of TUNEL staining of contralateral hippocampi at 3 days post-CCI. (B-C) TUNEL positive cells in (B) ipsilateral and (C) contralateral perilesional context of WT and *Nlrp3^-/-^* mice 3 days following CCI. (D-F TUNEL positive cells in (D) ipsilateral and (F) contralateral perilesional hippocampi of WT and *Nlrp3^-/-^*mice 3 days following CCI. Unpaired t-test. n = 6 secctions per mice from 3-4 mice per genotype and time point. Individual data points biological mean of cells per section. All data are presented as mean ± SEM.

**Figure S2.**
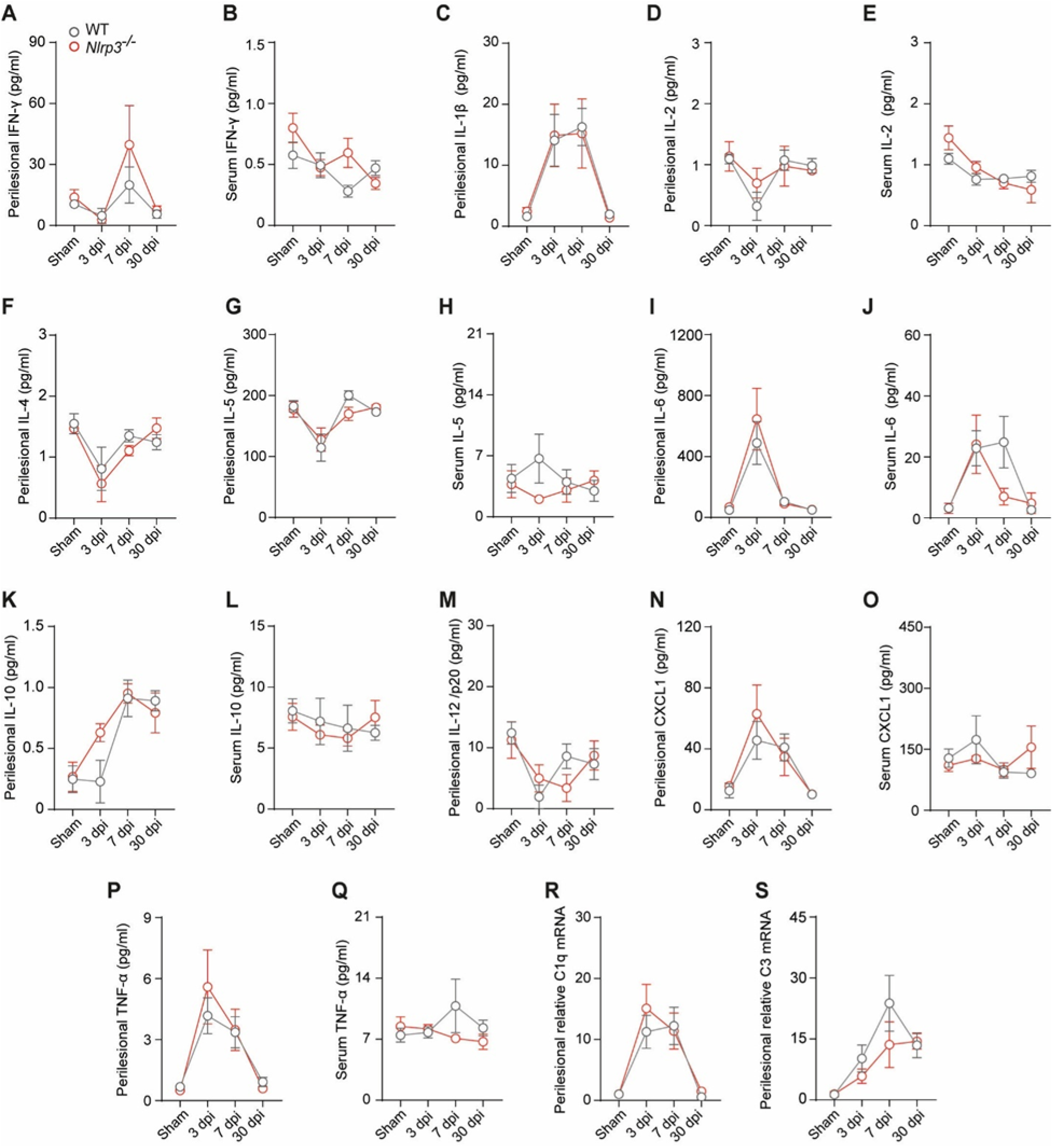
Key pro-inflammatory markers are regulated for up to 30 days in perilesional tissue and serum following CCI. (A–Q) Inflammatory cytokine levels in perilesional tissue and serum of WT and *Nlrp3^-/-^* mice following CCI. (A) Increased perilesional IFN-γ at 7 dpi in *Nlrp3^-/-^* mice. (B) Decreased serum IFN-γ at 30 dpi in *Nlrp3^-/-^* mice. (C) Upregulation of perilesional IL-1β at 3 and 7 dpi in both WT and *Nlrp3^-/-^* mice. (D) No significant changes in perilesional IL-2 in WT or *Nlrp3^-/-^* mice. (E) Decreased serum IL-2 at 7 and 30 dpi in *Nlrp3^-/-^* mice. (F) Decreased perilesional IL-4 at 7 dpi in *Nlrp3^-/-^* ^mice^. (G) Decreased perilesional IL-5 at 7 dpi in *Nlrp3^-/-^* ^mice^. (H) No significant changes in serum IL-5 in WT or *Nlrp3^-/-^* ^mice^. (I) Increased perilesional IL-6 at 3 dpi in both WT and Nlrp3-/-mice. (J) Increased serum IL-6 at 3 dpi in *Nlrp3^-/-^* and at 7 dpi in WT mice. (K) Increased perilesional IL-10 at 7 and 30 dpi in both WT and *Nlrp3^-/-^* mice. (L) No significant changes in serum IL-10 in WT or *Nlrp3^-/-^* mice. (M) Decreased perilesional IL-12 at 3 dpi in WT mice. (N) Upregulation of perilesional CXCL1 at 3 and 7 dpi in *Nlrp3^-/-^* mice. (O) No significant changes in serum CXCL1 in WT or *Nlrp3^-/-^* mice. (P) Upregulation of perilesional TNF-α at 3 dpi in *Nlrp3^-/-^*mice. (Q) No significant changes in serum TNF-α in WT or Nlrp3-/- mice. (R–S) Perilesional C1q and C3 mRNA levels. (R) Upregulation of perilesional C1q mRNA at 3 dpi in both WT and *Nlrp3^-/-^*mice. (S) Upregulation of perilesional C3 mRNA at 7 dpi in WT mice. n = 5 mice/genotype/timepoint. Two-way ANOVA. Data are presented as mean ± SEM. Post hoc analyses were performed using Tukey’s test.

**Figure S3.**
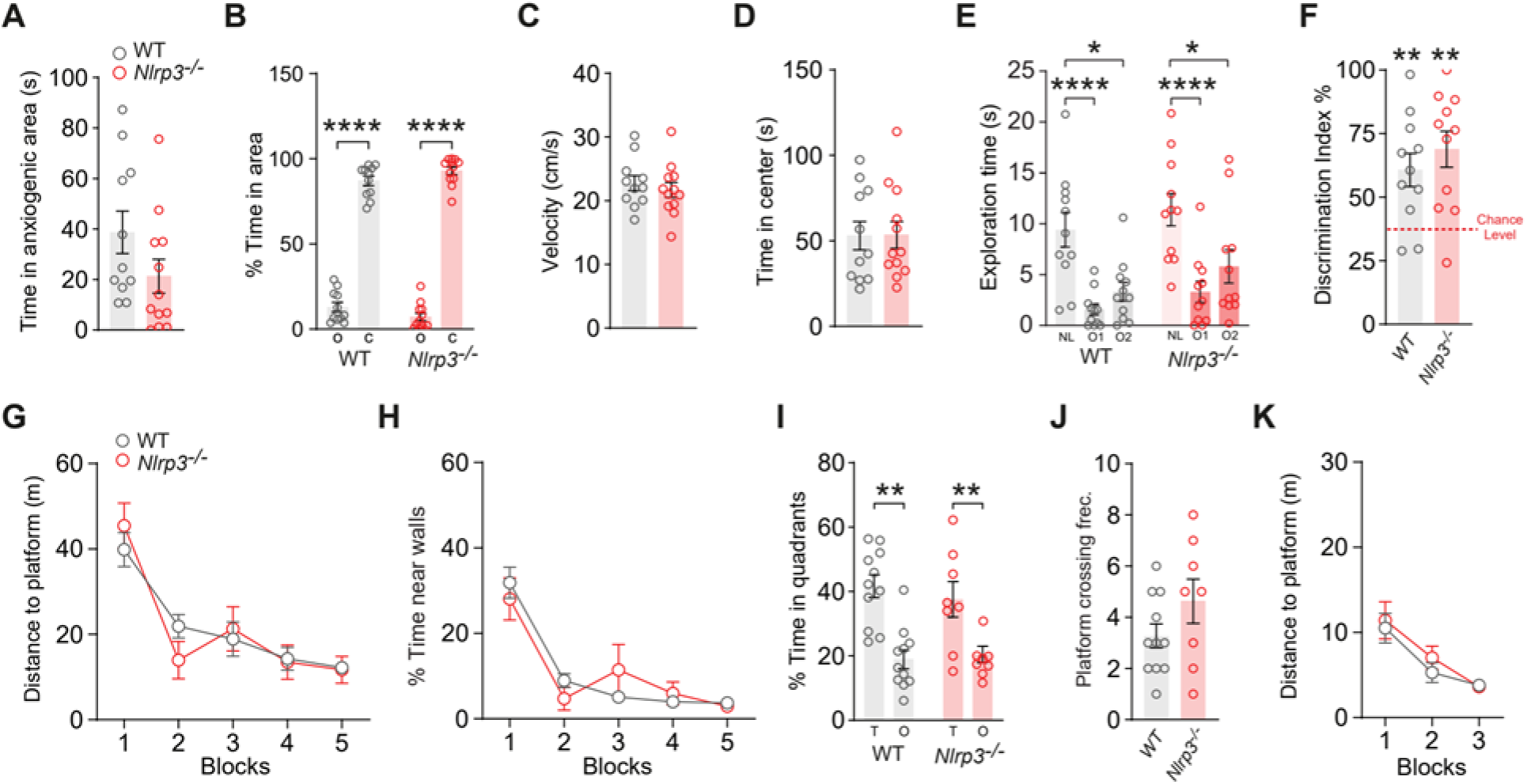
*Nlrp3^-/-^* mice show comparable behavior to WT controls. (A) *Nlrp3^-/-^* mice spent comparable time in anxiogenic areas compared to WT controls. n = 11-12 mice per genotype. Unpaired t-test. (B) Both genotypes spent significantly less time exploring the open sector than the closed sectors in the elevated zero maze. n = 11-12 mice per genotype. (C) Locomotor velocity was comparable between *Nlrp3^-/-^* and WT mice. n = 11-12 mice per genotype. Unpaired t-test. (D) Time spent exploring the center of the open field did not differ between *Nlrp3^-/-^* and WT mice. n = 11-12 mice per genotype. Unpaired t-test. (E) *Nlrp3^-/-^* mice displayed intact short-term novel location recognition memory, as indicated by significantly increased exploration of the object in a new location. n = 11-12 mice per genotype. (F) Both WT and *Nlrp3^-/-^* mice showed a significant discrimination index above chance. n = 11-12 mice pe genotype. One-sample t-test. (G-K) *Nlrp3^-/-^* mice exhibited normal spatial learning in the Morris Water Maze, as shown by (G) a progressive decrease in distance to the platform. Two-way ANOVA with repeated measurements. (H) The percentage of time spent near the walls was similar between genotypes. Two-way ANOVA with repeated measurements. (I) Long-term spatial memory, assessed by quadrant occupancy, was comparable between *Nlrp3^-/-^* and WT controls. Two-way ANOVA. (J) as was the frequency of platform crossings 24 h after training. Unpaired t-test. (K) Cued learning performance was similar in *Nlrp3^-/-^* and WT controls, as indicated by a progressive reduction in distance to the cued platform Two-way ANOVA with repeated measurements. n = 8-11 mice per genotype. Data are presented as mean ± SEM.

**Figure S4.**
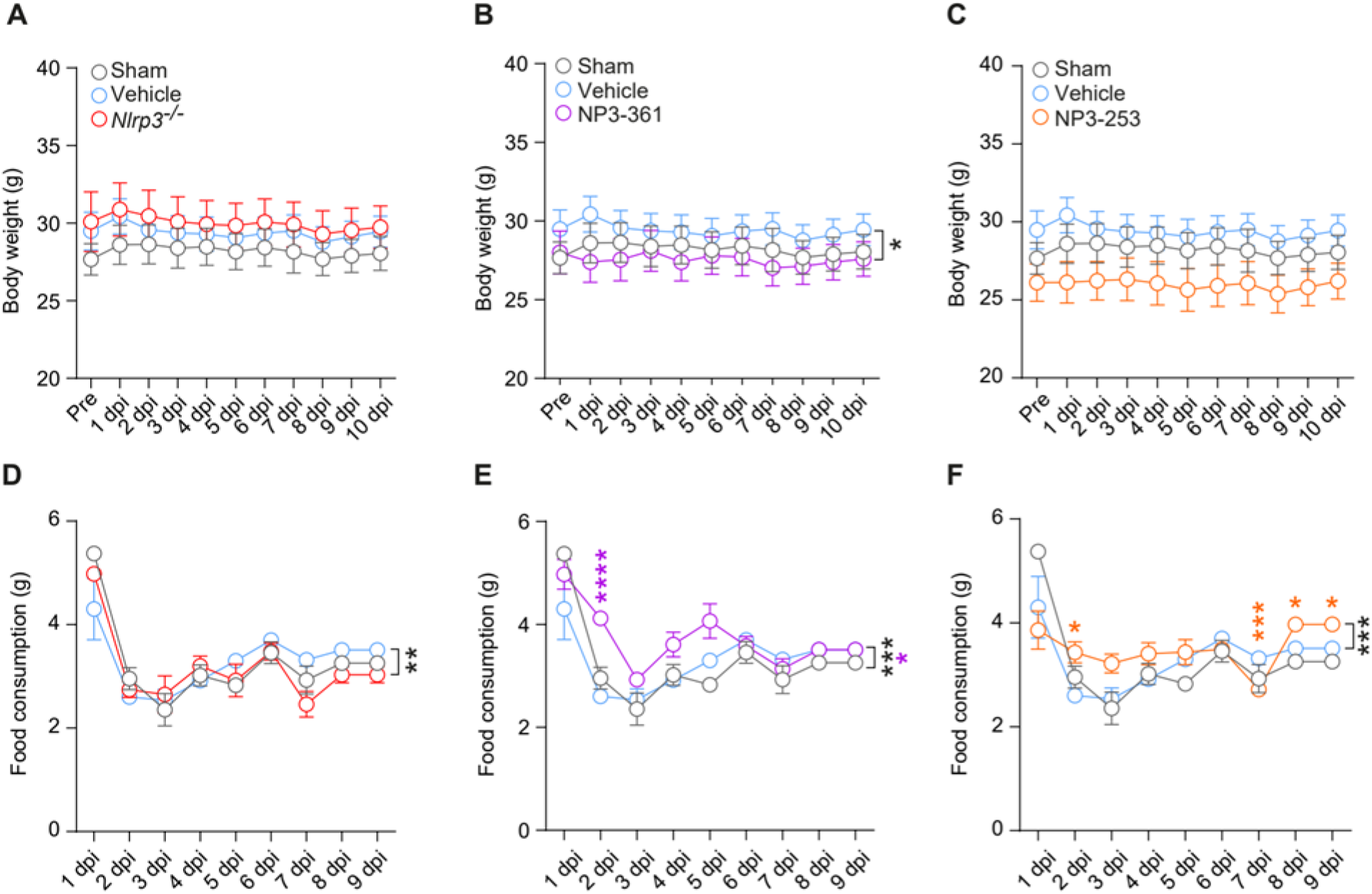
Body weight and food consumption during treatments after CCI. (A-C) Body weight during CCI and NLRP3 inhibitors treatment. (A) No significant changes in body weight following CCI in *Nlrp3^-/-^* mice compared to Sham and CCI injured WT mice. (B) Interaction effect of treatment and time in NP3-361 treated mice. Two-Way ANOVA Two-way ANOVA with repeated measures. (C) No significant changes in body weight following CCI in NP3-253 treated mice compared to Sham and CCI injured WT mice. (D-E) Estimated food consumption during CCI and NLRP3 inhibitors treatment. (D) Interaction effect of genotype and time in *Nlrp3^-/-^*. (E) NP3-361 and (F) NP3-253 treated mice consume significantly more food containing inhibitors during treatment following CCI. n=10-11 mice per treatment / genotype. Two-Way ANOVA Two-way ANOVA with repeated measures. All data are presented as mean ± SEM. Significance is indicated in color for genotype/treatment effects and in black for interaction effects. *p < 0.05, **p < 0.01, ***p < 0.001, ****p < 0.0001.

