## Supplementary figures and images for "NLRP3 inhibition maintains microglia architecture and enhances behavioral recovery after traumatic brain injury"

### Figure S1

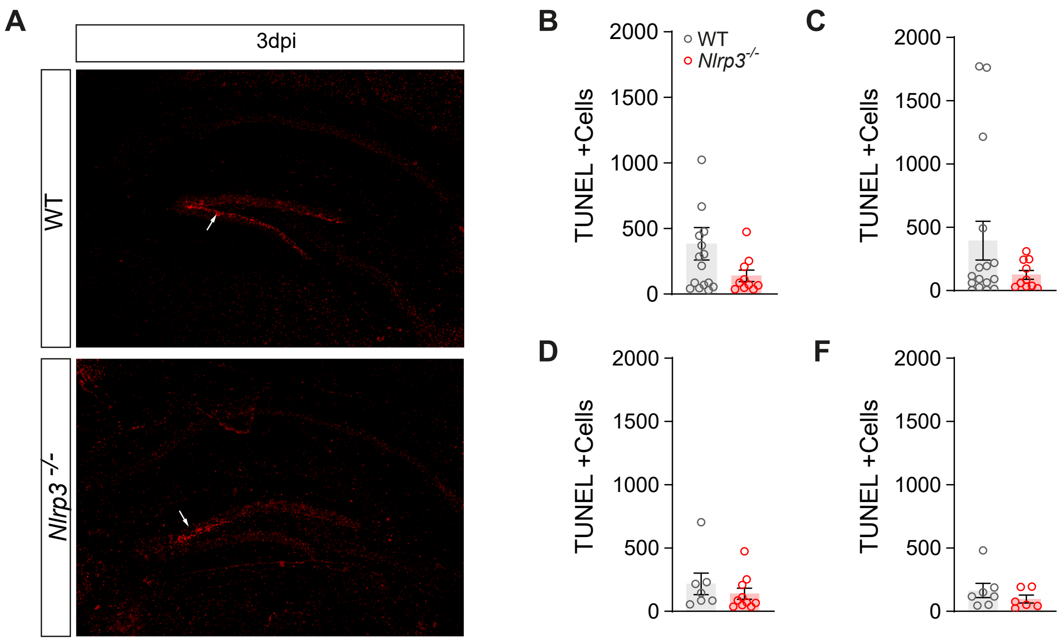

### Figure S2

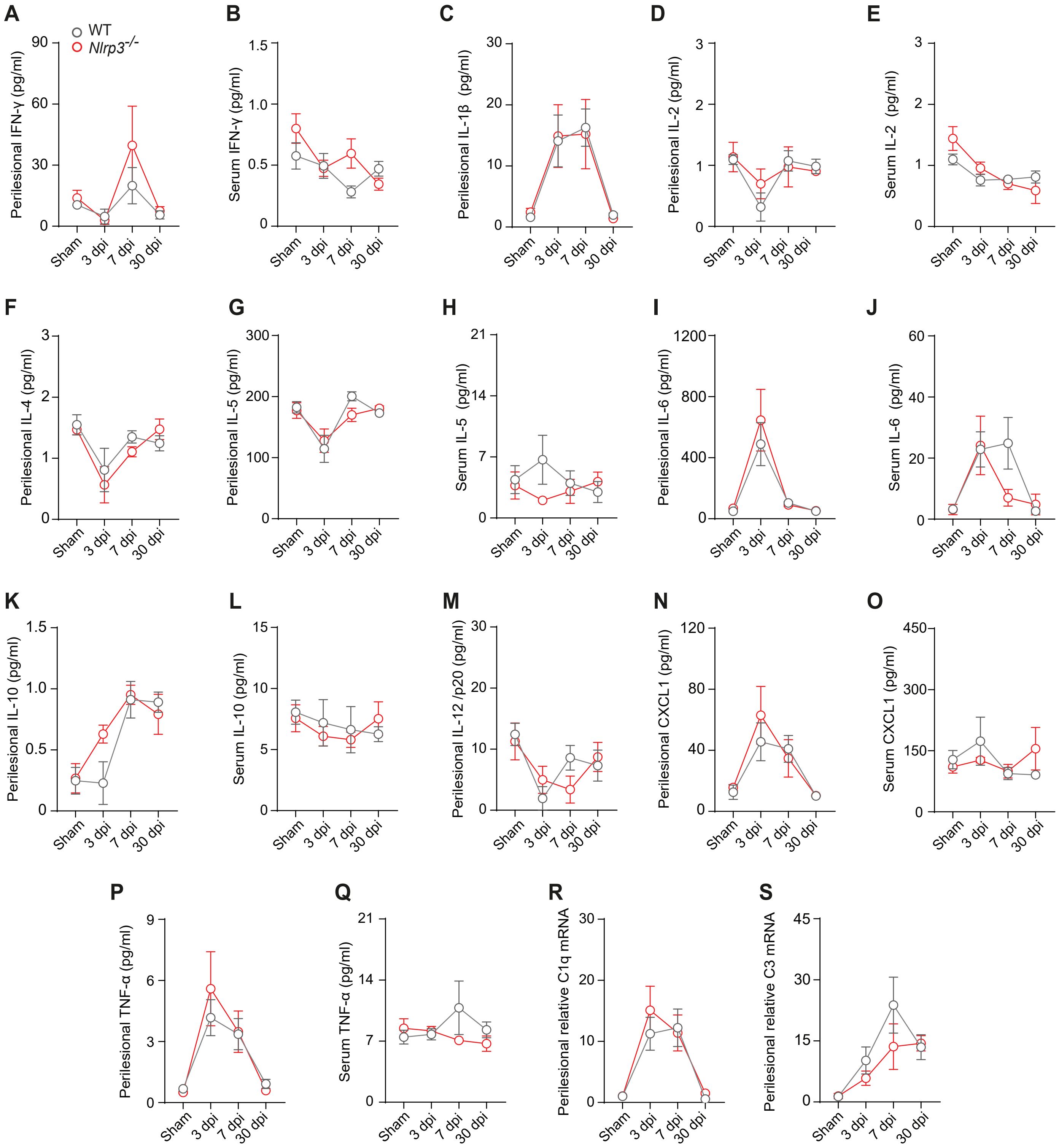

### Figure S3

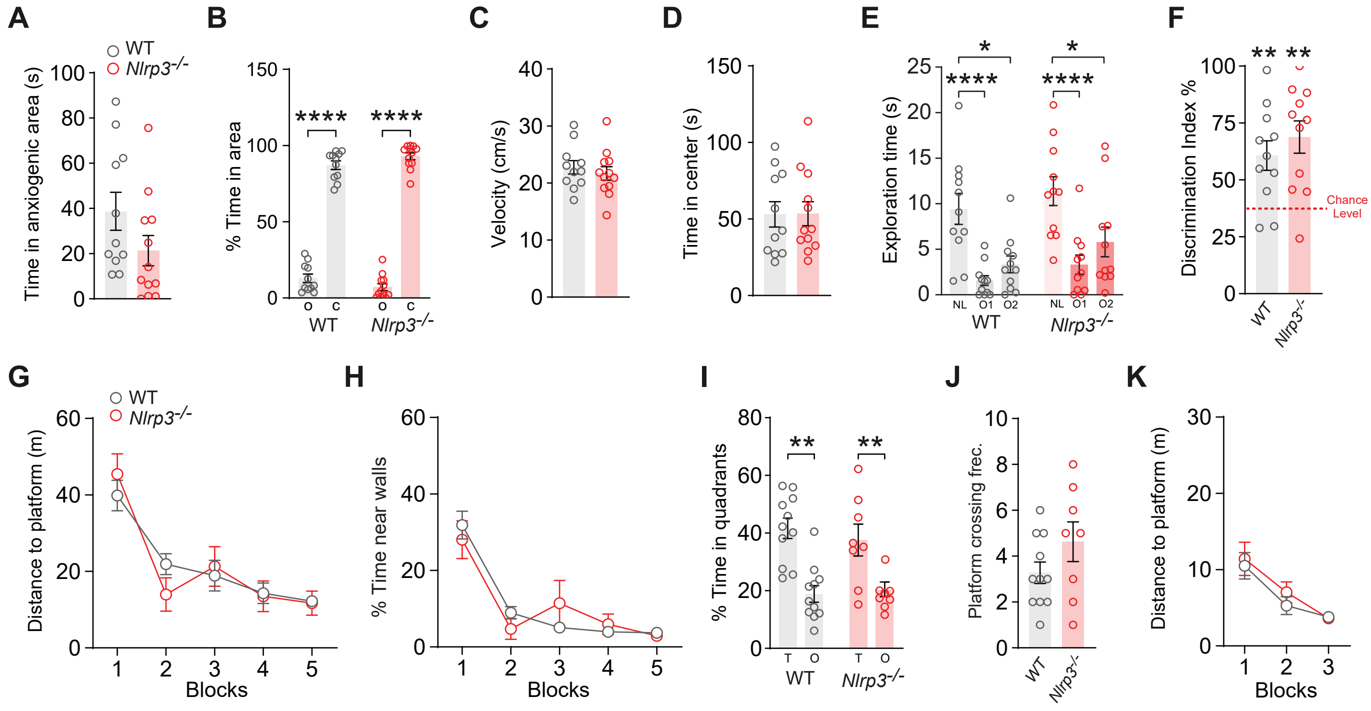

### Figure S4

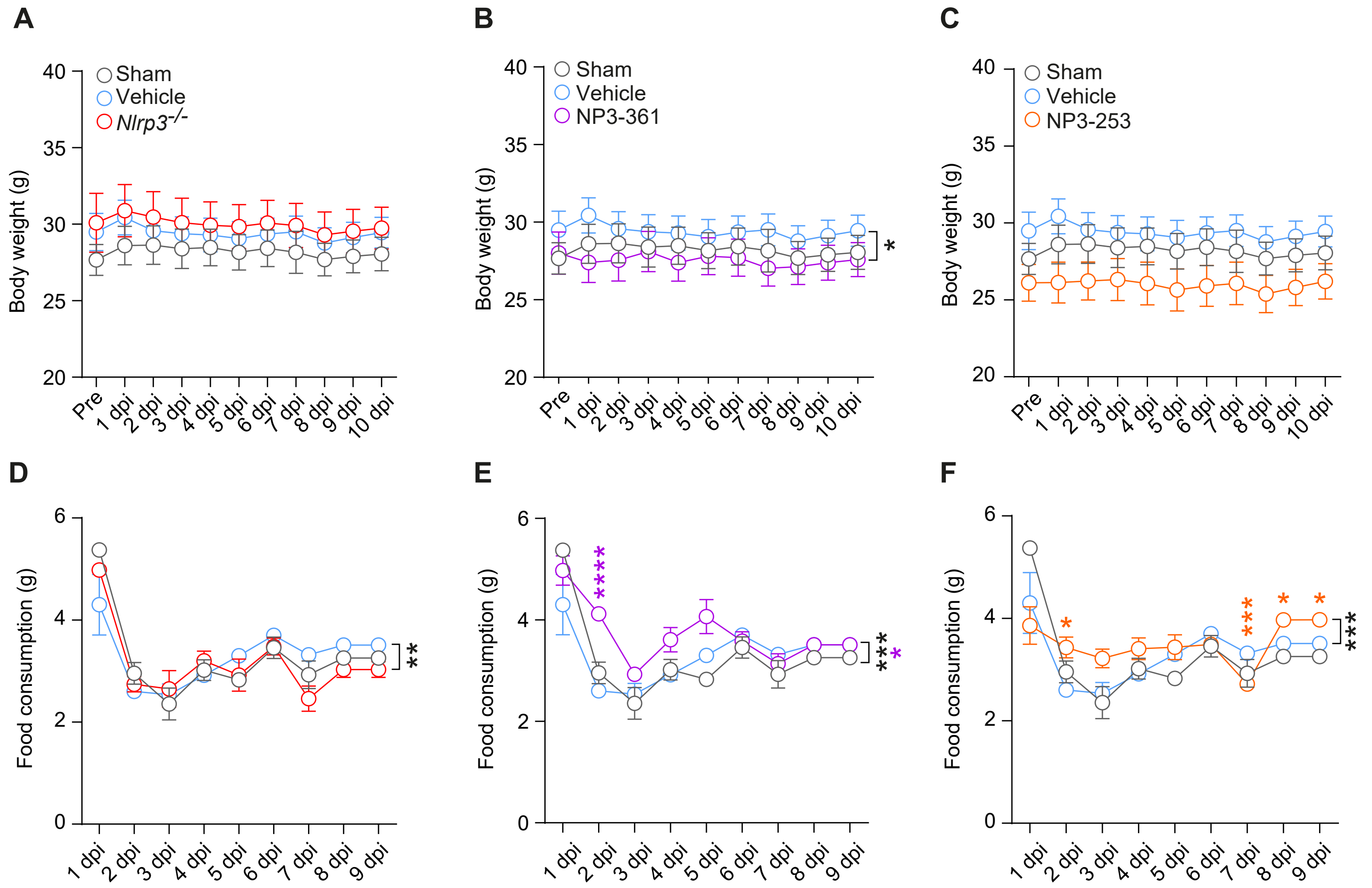
